# Offsetting ROS-mediated arrest of endothelial fenestration dynamics permits long-term optical super-resolution- and AFM-imaging

**DOI:** 10.1101/2025.10.03.680369

**Authors:** Annika Kiel, Marcin Luty, Angela Kralemann-Köhler, Laureen Patricia Helweg, Jasmin Schürstedt-Seher, Jerzy Kotlinowski, Jakub Pospíšil, Malgorzata Lekka, Thanh-Diep Ly, Thomas Huser, Jan Schulte am Esch, Wolfgang Hübner, Karolina Szafranska, Bartlomiej Zapotoczny

## Abstract

Advances in cell biology demand methods that resolve the structure and dynamics of subcellular organelles in living cells. Live-cell super-resolution fluorescence microscopy meets this need but is constrained by phototoxicity, which disturbs cellular function and biases interpretation. Liver sinusoidal endothelial cells (LSECs), with their physiologically critical and highly dynamic fenestrations, represent a particularly challenging model system. We show that photoactivation-generated reactive oxygen species (ROS) are the primary cause of fenestration arrest during fluorescence imaging. Using three-dimensional structured illumination microscopy (3D SR-SIM), we systematically evaluated fluorophores and ROS scavengers to optimize imaging conditions. A combination of BioTracker staining and CO₂-independent medium supplemented with N-acetylcysteine (NAC) preserved fenestration dynamics without altering fenestration number or size. Complementary atomic force microscopy (AFM) confirmed ROS-dependent impairment of fenestration dynamics and revealed nanomechanical changes upon illumination. These findings establish the mechanism underlying imaging-induced artefacts in LSECs and provide a broadly applicable strategy to extend live-cell super-resolution microscopy.

## 1. Introduction

New advanced imaging approaches, capable of capturing dynamic processes in living cells, are needed to continue our growing understanding of subcellular structures and their molecular mechanisms^1,2^. Tracking dynamic processes at the nanoscale requires not only high spatial and temporal resolution but also non-invasive image acquisition to preserve cell viability and fluorophore stability. Transcellular *fenestrations* in liver sinusoidal endothelial cells (LSECs) represent one of the most fragile and difficult cellular features to be visualized^3^. These structures regulate passive size-dependent filtration of blood from plasma proteins (e.g., albumin), smaller lipoproteins such as chylomicron remnants, VLDL particles, viruses or drug molecules, while excluding larger particles such as intact chylomicrons, blood cells, and other macromolecular debris. Fenestration size (50-350 nm), dynamics (opening/closing in seconds) and delicate nature (easily influenced by many factors) pose particular challenges for imaging, especially in living cells^4,5^.

The dynamic character of fenestration was proposed already in 1995^6^ but only revealed in 2017 using the latest advances in atomic force microscopy (AFM)^7^. This label-free technique allowed for studying fenestration in genetic models^8^, under the influence of various agents^7,9^ as well as assessing their dynamic behavior *in vitro*. Fenestrations have been shown to continuously open, close, change diameter up to 200% and migrate several micrometers, all within their short average lifespan of ∼20 minutes^10^. Although AFM presented tremendous potential in tracking fenestration dynamics, the trade-off between the field of view size and spatial/temporal resolution, together with the lack of chemical information necessitates the development of alternative techniques.

Today’s knowledge of LSEC ultrastructure is based on live cell studies with AFM and fixed fluorescently labelled cells imaging using nanoscopy techniques such as direct stochastic optical reconstruction microscopy (dSTORM)^9^, stimulated emission depletion microscopy (STED)^11^ and super-resolution structured illumination microscopy (3D SR-SIM)^11,12^. There is a growing demand for super-resolution imaging strategies capable of resolving fine cellular architecture while being suitable for live-cell, long-term observations. 3D SR-SIM addresses this need by enabling acquisition of large fields of view within relatively short imaging times and <200 nm resolution. Although 3D SR-SIM typically achieves about a twofold improvement in spatial resolution compared to conventional wide-field microscopy — less than other super-resolution techniques — it offers several key advantages. Its compatibility with standard wide-field microscope configurations, wide selection of compatible fluorophores, and relatively low laser intensity make it particularly suitable for biological applications. These characteristics, combined with its large field of view capability, make 3D SR-SIM particularly promising for long-term live cell imaging of LSECs^13–15^.

Nevertheless, existing reports on the nanoscopy applications for studying LSEC present very limited fenestration dynamics with attenuated responsiveness to stimuli such as oxidized low-density lipoprotein (oxLDL)^16^ or the actin depolymerization agent, cytochalasin D^17,18^. These attempts at studying fenestration response were hampered by fenestrations losing their dynamics and appearing arrested after labelling and illumination. In particular, Martino and coworkers highlighted phototoxicity as a potential factor preventing prolonged imaging of fenestrations dynamics with STED. Another hurdle to prolonged live cell imaging is the limited photostability of fluorophores. Therefore, many recent reports have focused on the creation and synthesis of novel fluorescent proteins and organic fluorophores with significantly improved photostability^19–22^. Improving photostability only partly solves the problem, because the phototoxic effects of visible light on cells must also be addressed.

In this study, we optimized a SIM-based approach for prolonged imaging of fenestration dynamics. We use LSEC fenestrations as a nanoscale, easily quantifiable model (with parameters such as number, diameter, lifespan, motility, deformability, and stiffness) to optimize super-resolution imaging conditions that generate insights widely applicable to live-cell imaging strategies of other cell types. By comparing label-free AFM data with corresponding SIM results, we confirmed that the observed phototoxicity is linked to light-induced reactive oxygen species (ROS) in the presence of the fluorescent dye. To mitigate excess ROS formation and preserve fenestration dynamics, we successfully implemented media supplementation with N-acetyl-L-cysteine (NAC). This strategy can be widely implemented for other optical techniques used for live cell imaging, possibly revealing previously omitted dynamic events due to phototoxicity. Overall, the proposed method enables prolonged live imaging with maintained cell dynamics, even in ROS-sensitive structures such as liver fenestrations.

## 2. Results

### 2.1 Selection of Fluorescent Dyes for Prolonged Live-Cell Super-Resolution Imaging of LSEC Fenestrations

Despite the low intensity of individual illuminations required for 3D SR-SIM imaging, membrane dyes suitable for live-cell super-resolution imaging must exhibit high photostability. To identify the most suitable dye for imaging LSEC fenestrations, we tested several commercially available membrane stains, such as CellMask Plasma Membrane Stain Orange, Vybrant DiI cell-labeling solution and BioTracker 555 Orange Cytoplasmic Membrane Dye. Additionally, we tested CellMask Green Actin Tracking Stain which visualizes fenestrae-associated cytoskeletal rings, serving as an indirect method for fenestration labeling. All dyes were tested under identical conditions using 3D SR-SIM to assess their photostability, signal-to-noise ratio, staining specificity, and suitability for extended time-lapse acquisition. Live-cell imaging was performed over a total period of one hour, with images acquired every 15 minutes. Representative images for each dye are shown in **Figure 1**, with additional datasets provided in **Figures S1–S4**. A noticeable decrease in fluorescence intensity (photobleaching) was observed on almost all images for the CellMask membrane and actin dyes, after just a single acquisition (**Figure 1**, 15 min CellMask Actin, CellMask Membrane and **Figure S1 and S2**). For the Vybrant DiI dye, the combination of unspecific dye accumulation and labeling of cellular debris together with inefficient uptake by the cells, results in an exceptionally high intensity difference compared to the less densely labeled plasma membranes within the sieve plates and fenestrations (see **Figure 1**, Vybrant DiI, yellow arrowheads and **Figure S3)**. This remarkable dynamic intensity range poses significant challenges to efficient reconstruction of SIM images. The potential spherical aberrations caused by the bright dots are accentuated, thereby obscuring the lower intensity regions with sieve plates. In contrast, BioTracker provided consistent and strong staining across nearly all cells, with clearly visible plasma membrane architecture, high-contrast visualization of fenestrations and no noticeable photobleaching over the one-hour imaging period (**Figure 1**, BioTracker and **Figure S4**). Based on these dye screening results, BioTracker was selected for all further experiments.

**Figure 1.**
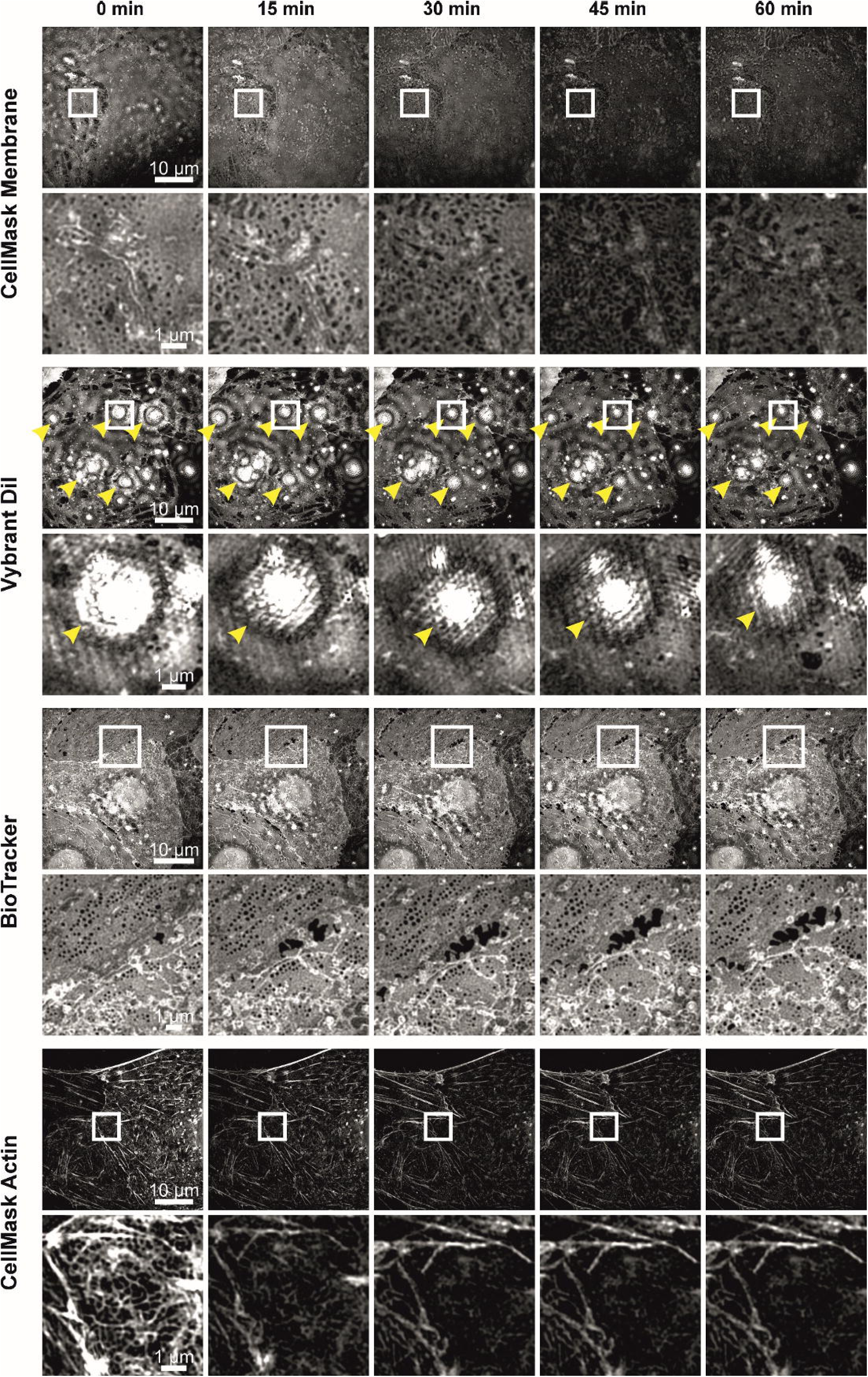
Live-cell 3D SR-SIM imaging of LSECs over a 1-hour time span using different membrane and cytoskeletal dyes. Representative time-lapse images of primary LSECs acquired with 3D SR-SIM are shown for four different dyes: CellMask Plasma Membrane Stain Orange, Vybrant DiI cell-labeling Solution, BioTracker 555 Orange Cytoplasmic Membrane Dye, and CellMask Green Actin Tracking Stain. Each panel displays selected LSEC imaged over the course of one hour in 15 minute intervals. For every condition, an overview of nearly the entire cell is shown together with a corresponding region of interest (ROI – white box) highlighting fenestration morphology and membrane dynamics over time. Yellow arrowheads indicate artifacts. In addition, dye performance over time was assessed, including photostability, signal-to-noise ratio and staining specificity.

### 2.2 Strategies to attenuate fluorophore-induced mitochondrial ROS release in live LSEC imaging

To identify the phototoxic effect of fluorophore activation during live-cell imaging of LSECs, intercellular ROS were quantified. In parallel, the potential protective effects of media supplementation with oxygen scavenger Oxyrase or the antioxidant N-acetylcysteine (NAC) were evaluated. An ATP-based viability assay demonstrated that neither Oxyrase nor NAC affects LSECs *in vitro* (**Figure 2A**). Based on our previous findings^12^, NAC was tested at concentrations between 0.5 to 2.0 mg/mL, while Oxyrase was applied at 1%, as recommended by the manufacturer.

**Figure 2.**
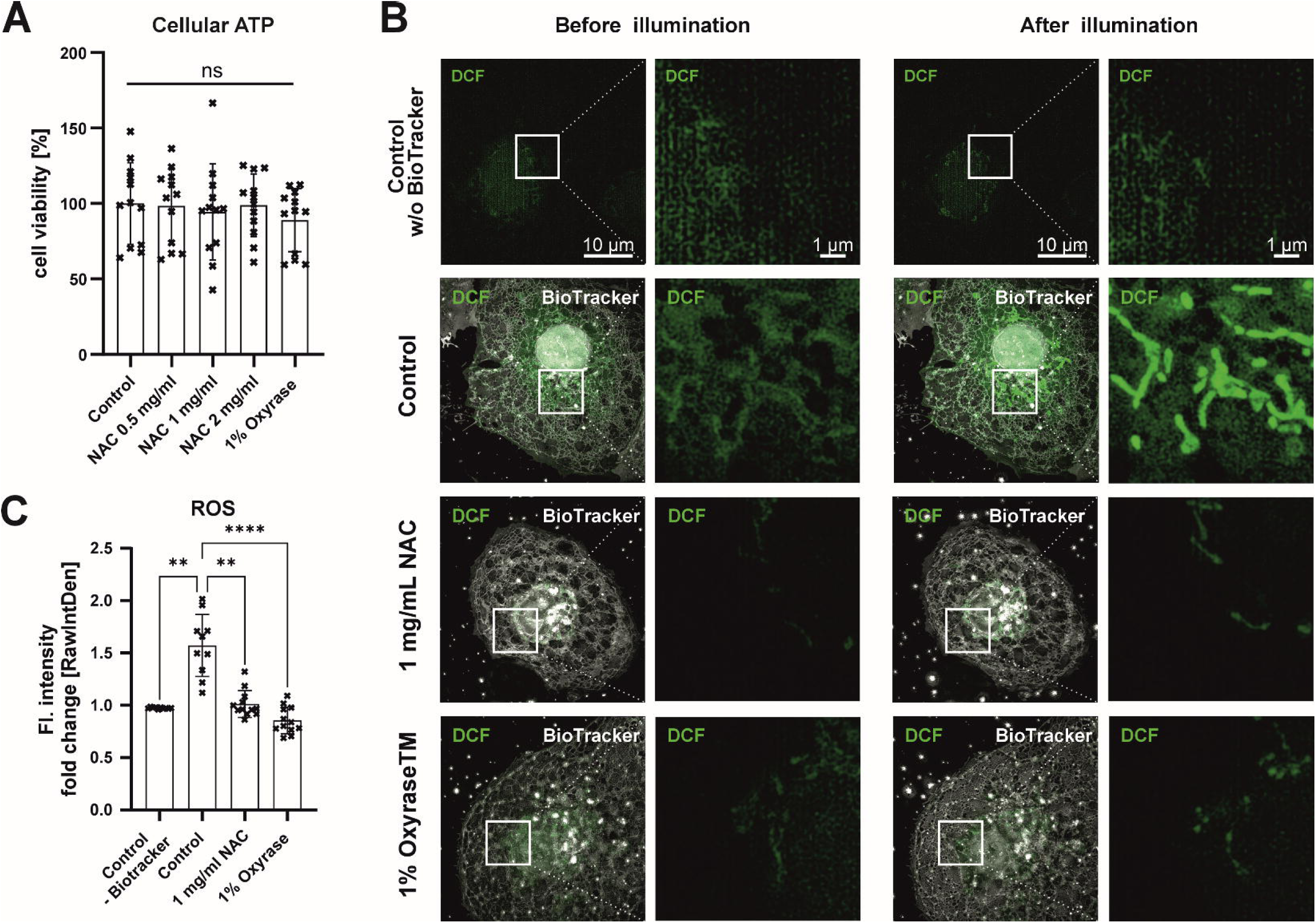
Light-induced intracellular ROS detection. **A)** The influence of different concentrations of N-acetylcysteine (NAC) and Oxyrase on cell viability was tested by measuring cellular ATP levels. Data were normalized to the untreated control. Mean ± SD. **B)** To detect intracellular ROS during live-cell imaging, LSECs were pre-treated with ROS indicator (DCFDA/H2DCFDA). Intracellular ROS levels were assessed before and after constant illumination every 3 min for a total of 45 min. The influence of the fluorescent dye BioTracker 555 itself on intracellular ROS levels was evaluated (Control w/o BioTracker 555 and Control), along with the effects of 1 mg/mL NAC and 1% Oxyrase. ROS activity was visualized by 3D SR-SIM imaging of the fluorescent signal resulting from the oxidation of DCFDA to fluorescent DCF. **C)** Fluorescence intensity was measured for at least 10 cells per treatment using ImageJ. The sum of fluorescence intensity after illumination was normalized to the intensity before illumination. Mean ± SD, each point represents the same field of view from the start to the end of the experiment, one-way ANOVA (Kruskal-Wallis test) with Dunn’s multiple comparison test, ** p < 0.001 and **** p < 0.0001.

Next, we investigated ROS formation and the impact of medium supplementation during 3D SR-SIM imaging. After preloading with DCFDA/H2DCFDA, a fluorescent ROS indicator, cells were illuminated every 3 minutes over a period of 45 minutes, and representative first and last frames are shown in **Figure 2B**. No increase in intracellular ROS was detected in the unstained controls. In contrast, BioTracker-stained cells in non-supplemented media showed clear ROS induction, predominantly localized to mitochondria in the perinuclear region. Supplementation with either 1 mg/ml NAC or 1% Oxyrase markedly reduced ROS formation during imaging. The quantitative analysis of the DCF fluorescent signal (**Figure 2C**) confirmed that both NAC and Oxyrase significantly suppressed light-induced ROS formation in LSECs, reaching levels comparable to the unstained controls. Overall, these results indicate that the observed phototoxic effect is mediated by ROS induced by the combination of the fluorophore and illumination.

### 2.3 Dynamics of fenestrations – 3D SR-SIM

To establish conditions suitable for prolonged 3D SR-SIM imaging of cellular dynamics, including tracking of LSEC fenestrations with minimal phototoxicity, we established an efficient live-cell imaging protocol (see *Materials and Methods* section, point 2.4.3). In non-supplemented media (**Figure 3A**, control), most of the cells became completely arrested, as indicated by loss of fenestration dynamics (**Figure 3A**, yellow circle) and absence of cell movement across time frames. Moreover, fenestration closure and membrane rupture were observed in other cells, indicating severe toxicity (**Figure S5**). Supplementation with NAC preserved cellular dynamics, as shown by sustained fenestration mobility (**Figure 3A**, green circles and green arrowheads) and active membrane remodeling and repair, including gap closure over time (**Figure 3A**, green arrows, **Figure S6**). The quantification of fenestration number at the start (0 min) and the end (60 min) of the experiments revealed no significant difference in fold change of fenestration count in both control and NAC-supplemented cells (**Figure 3B**). This stability indicates that 3D SR-SIM imaging enables visualization of fenestrations in living cells without altering the overall number of fenestrations. However, despite preservation of the fenestration number, the analysis of fenestration dynamics by tracking the position of individual fenestrations over time (speed) showed a significant reduction in mobility for the non-supplemented controls (**Figure 3C**). These findings were further supported by polar plots of individual fenestrations trajectories, revealing limited fenestration displacement for the controls, whereas fenestrations in NAC-supplemented cells exhibited broader and more dispersed movement patterns (**Figure 3D**).

**Figure 3.**
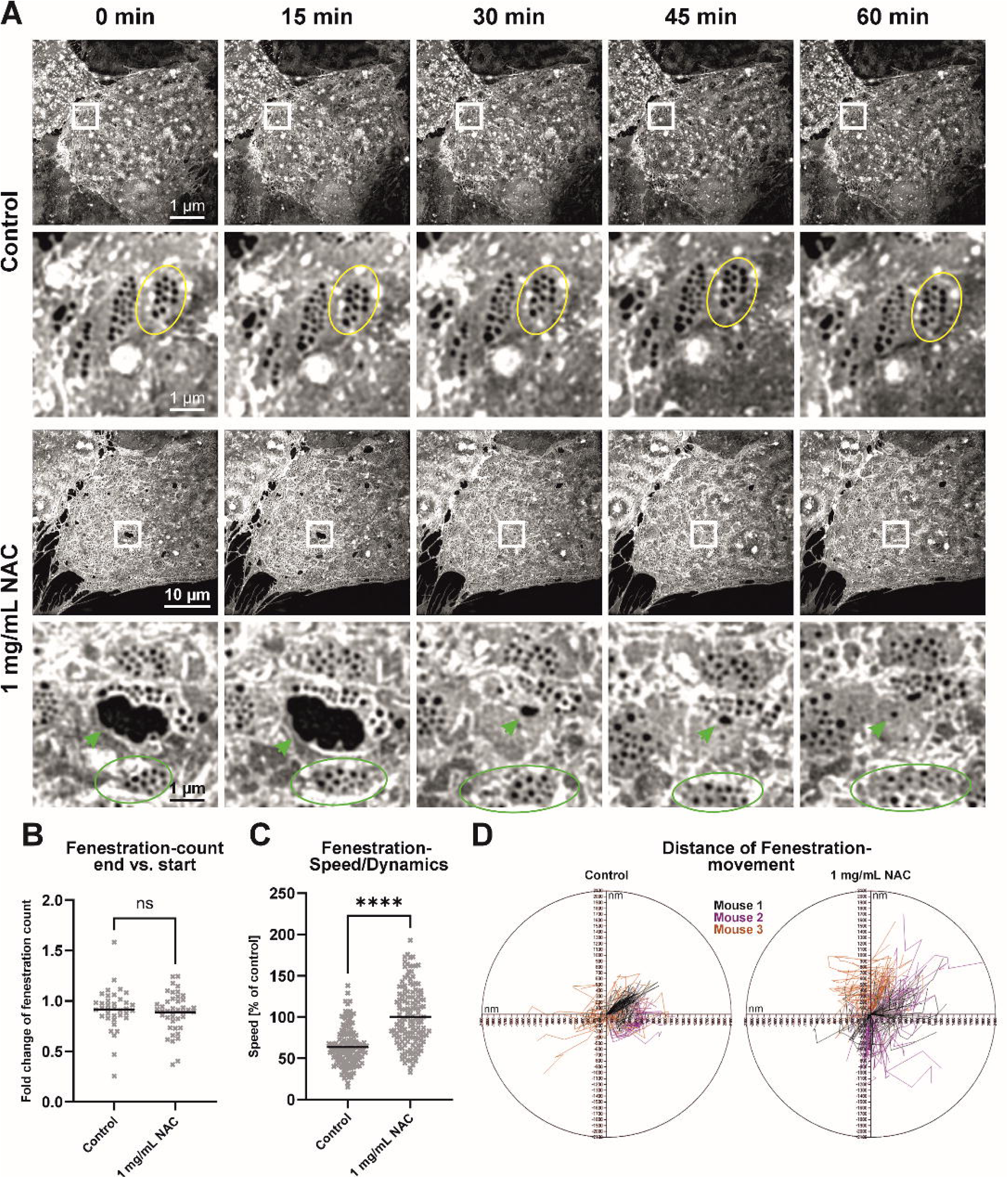
Effects of NAC supplementation on LSEC fenestration dynamics during 3D SR-SIM live-cell imaging. Cells were maintained at 37°C, stained with BioTracker 555, and kept in imaging medium. Time-lapse images were acquired every 15 minutes over a 60-minute period. **A)** Representative time-lapse images of LSECs without medium supplementation (control) or with 1 mg/mL NAC supplementation. In control cells, fenestrations remained arrested throughout the imaging period (yellow circles), with no observable membrane dynamics. NAC-supplemented cells exhibit preserved fenestration mobility (green circles) and active membrane remodeling (green arrowheads). Scale bars: overview panels, 10 µm; Region of interest, 1 µm. **B)** Fold change in fenestration count between start and end of the experiment was calculated (60 min / 0 min). No significant (ns) difference was observed between the groups. Each dot represents the same field of view for the start and end of the experiment. Mean + datapoints; n = 3 animals, unpaired Mann-Whitney rank test. **C)** Quantification of fenestration dynamics, expressed as displacement between time points (speed). NAC treatment significantly increased fenestration mobility relative to control. Mean + single datapoints, n = 3 animals, unpaired Mann-Whitney rank test, ****p < 0.0001. **D)** Polar plots showing individual fenestration displacement from three biological replicates (mice). NAC-treated cells show increased movement relative to the non-supplemented control.

Similarly to NAC, Oxyrase was evaluated as a supplement for live-cell imaging using 3D SR-SIM (**Figure S7** and **S8**). Although Oxyrase supplementation preserved the overall fenestration count (**Figure S9**), it failed to sustain essential indicators of cellular dynamics. In particular, no membrane remodeling activity or fenestration dynamics was observed in LSECs, which remained comparable to the non-supplemented control cells (**Figure S7** and **S8**). Consequently, only NAC supplementation was pursued for further evaluation.

### 2.4 Quantitative evaluation of the effects of fluorophore and illumination conditions on fenestration dynamics using AFM

#### 2.4.1 BioTracker

To further evaluate potential perturbations of fenestration dynamics by fluorescence microscopy, we used label-free AFM imaging to separate light-induced from dye-induced (chemical) contributions to the phototoxic effect and focus on mitigation strategies. By maintaining continuous AFM imaging over three hours, we sequentially altered the extracellular environment in four steps: (1) control imaging medium, (2) imaging medium containing fluorophore, (3) fluorophore washout, and (4) subsequent exposure to fluorescence illumination. NAC was included as a supplement in all steps of the experiment. A representative experiment for BioTracker is presented in **Figure 4**, where 15 images from a total of 95 images were shown (**Figure 4C**; **Video S1-S2**). Quantification of fenestration speed (**Figure 4B, D**) showed that labelling with BioTracker alone did not impair fenestration dynamics. Supplementation with 1 mg/mL NAC preserved fenestration mobility across all the steps of the experiment, although a modest (∼25%) decline was detected after the second illumination. In AFM experiments, the protective effect of Oxyrase used as a supplement was not observed (**Figure S10, Video S3**). Gap formation (>350 nm) limited its suitability for long-term AFM imaging, and the motility of remaining fenestrations was ∼50% slower than before illumination. Parallel experiments were performed for BioTracker without supplementation (**Figure S11**). About 50% reduction in fenestration motility was observed, most fenestrations becoming arrested in minutes after illumination with only minor collective cell movement (**Figure S11, Video S4**).

**Figure 4.**
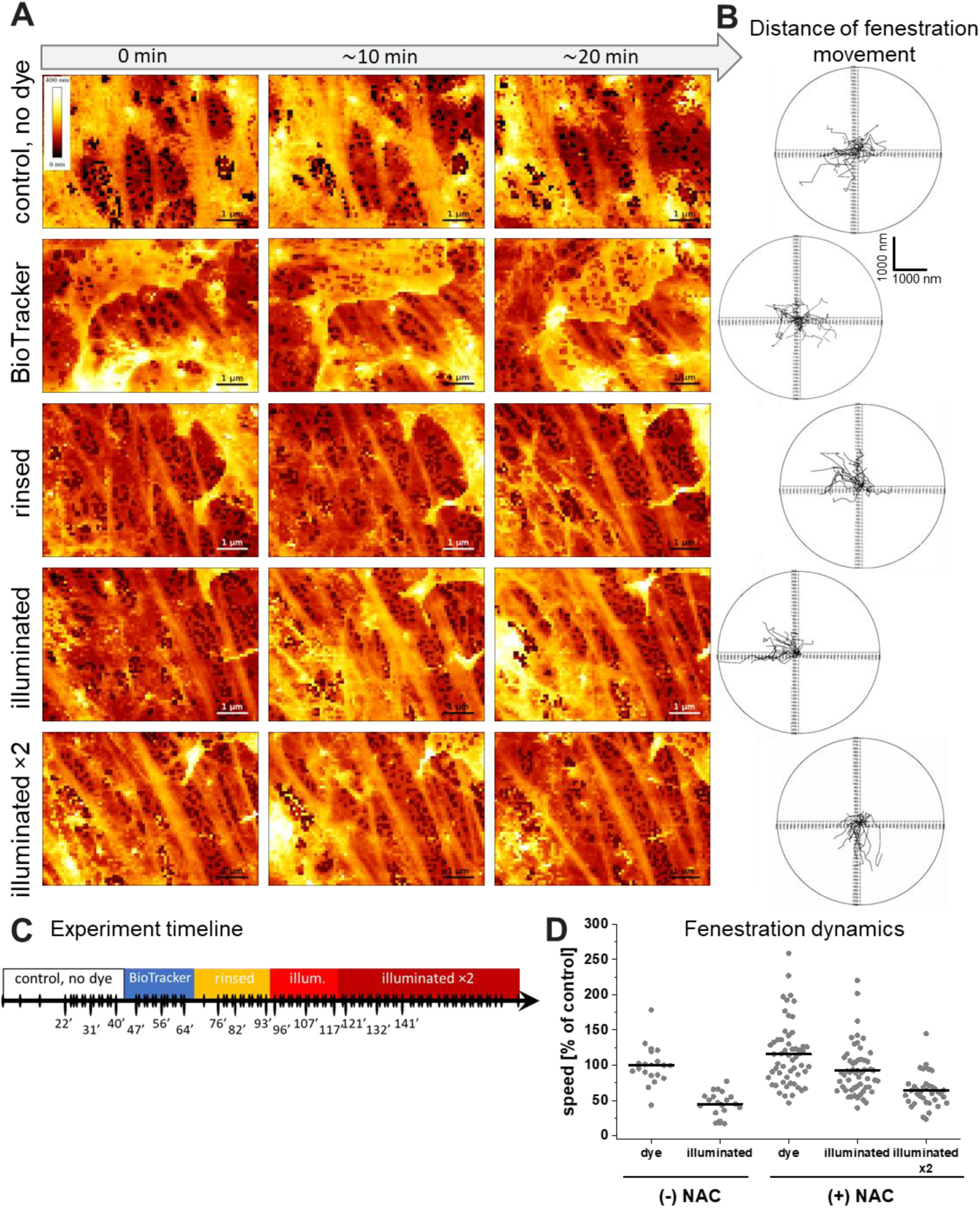
Effects of NAC supplementation on LSEC fenestration dynamics during AFM live-cell imaging. At each step, LSECs were measured in imaging medium supplemented with 1.0 mg/ml NAC at 37°C. **A)** Representative time-lapse images from steps 1-4 (rows 1-4). **B)** Polar plots of individual fenestration displacement at the corresponding steps. **C)** Graphical representation of time-lapse imaging: continuous small-area scans were acquired approximately every 2 minutes for 3 h, interweaved with large-area scans lasting ∼8 minutes, performed as needed to verify cell condition. Points labeled with timestamps correspond to the images shown in **(A)**; the remaining frames are available in **Video S1**. **D)** Quantification of fenestration speed indicates that two illumination events preserve fenestration dynamics more effectively than a single illumination event in the absence of NAC supplementation.

#### 2.4.2 Other dyes

Similarly to BioTracker, neither CellMask Membrane nor CellMask Actin dyes altered fenestration dynamics in the absence of illumination, suggesting that it is not the dye alone that induces the observed changes. The effect of illumination on fenestrations was particularly pronounced in cells labeled with CellMask Membrane dye without media supplementation. Fenestrations became arrested within seconds after light exposure (**Figure S12, Video S4**). In addition, illumination induced a transient enlargement of fenestration diameter, which returned to baseline within a few minutes. This photosensitive response was reversible and reproducible, as repeated illumination triggered a comparable, temporary increase in fenestration size. Interestingly, for CellMask Actin dynamics of the fenestration-associated cytoskeleton remained unchanged even after the second illumination (**Figure S13**). This indicates that the approach of indirect labeling of actin can provide an effective strategy to track fenestrations, if bleaching is mitigated.

To further validate our observations that ROS generation directly impairs fenestration dynamics, we reanalyzed our previously published datasets, in which hydrogen peroxide (H_2_O_2_) was used as a direct, well-established intracellular ROS inducer (see ^12^; experimental context in **Figure 4** of that study). Applying the current analytical approach to quantify fenestration dynamics, we confirmed that the treatment with 50 μM H_2_O_2_ led to an over 80% reduction in mean fenestration speed (**Figure S14**), consistent with the observations reported here during fluorophore photoactivation.

#### 2.4.3 Cell nanomechanics in experiments with BioTracker

Finally, we evaluated changes in cell nanomechanics, which are closely linked to fenestration regulation via the cytoskeleton. AFM was used to measure and calculate the apparent Young’s modulus of LSECs (**Figure 5**). No changes in apparent Young’s modulus were detected until the fluorescent illumination, when significant cell softening was observed (**Figure 5A**). Control experiments (mock, without fluorophore) showed no changes in cell mechanics in culture over time, even after illumination (**Figure 5B**). The results indicate that cytoskeletal remodeling occurs after photoactivation of fluorophores, demonstrating that both the fluorophore and its activation are required to induce the observed changes. Taken together, the nanomechanical findings are in line with the study of fenestration dynamics, verifying that the negative effect of fluorescence imaging can be attenuated by NAC supplementation, although not to a full extent.

**Figure 5.**
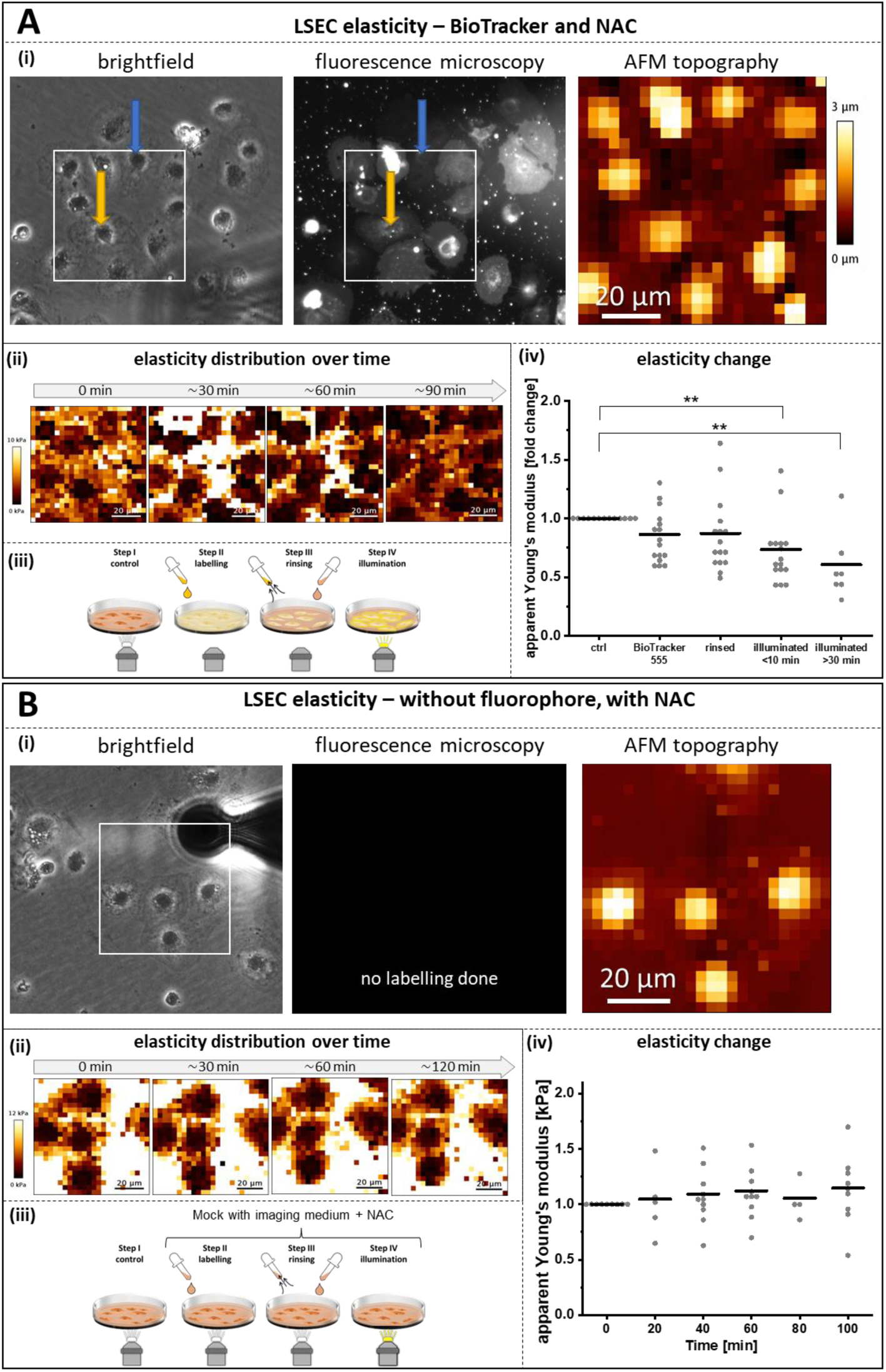
Effects of BioTracker and photoactivation on LSEC nanomechanics during AFM live-cell measurements. **A**) LSECs labelled with BioTracker and imaged in NAC-supplemented medium at 37⁰C. (i) Representative brightfield image from imaging step 1 and fluorescence microscopy image from imaging step 4. Orange and blue arrows indicate cells with different labelling intensities. The white box marks the area analyzed by AFM; with the corresponding AFM topography channel shown. (ii) Apparent Young’s modulus maps of the cells measured at steps 1-4 of the experimental setup. (iii) Schematic overview of the experimental workflow; steps 2 and 3 were performed without illumination. (iv). Apparent Young’s modulus measured sequentially in the central region of the same selected cells. Each data point represents an individual LSEC. Data obtained from three independent biological replicates. ** p < 0.01 **B**) Control experiment without fluorophore (mock treatment). (i) Representative brightfield image of cells selected for AFM analysis and a representative fluorescence image showing no signal due to the absence of fluorophore labeling. The white box indicates the AFM measurement area, with the corresponding AFM topography channel shown. (ii) Apparent Young’s modulus maps of the cells measured at different timepoints (iii) Schematic overview of the experimental workflow; all imaging steps performed under dim white light – no fluorescence excitation or illumination was used. (iv) Apparent Young’s modulus measured in the central region of the same selected cells. Each data point represents an individual LSEC. Data were obtained from two independent biological replicates. No significant changes in cell elasticity were detected over time.

## 3. Discussion

Phototoxicity has long been a fundamental challenge limiting the application of super-resolution optical techniques in live cell imaging. In this study, we established a protocol for prolonged live-cell imaging with 3D SR-SIM by combining BioTracker staining with NAC supplementation and demonstrated its applicability for studying the dynamic behavior of liver fenestrations.

For studying the exact mechanisms of phototoxicity, we incorporated AFM, a label-free technique, with 3D SR-SIM. AFM imaging and force spectroscopy enabled separation of light- and dye-induced effects, providing mechanistic insights into the impairment of fenestration dynamics upon illumination. As a result, we provided evidence that previous reports attempting to implement live-cell imaging of LSEC fenestrations had severe limitations, mainly due to phototoxicity, resulting from the use of incompatible fluorophores, lack of imaging media pH/CO_2_-balance and absence of ROS scavenging systems. In particular, CellMask Membrane dye (which has been successfully used for fixed cells) led to significant fenestration arrest, hampering dynamic observations in living cells of drug effects^16,23–25^. Martino and coworkers, (2018) used STED microscopy for LSECs stained with CellMask Orange and CellTracker Red and presented arrested fenestration dynamics, enlargement of fenestrations and restricted response to cytochalasin D (a known fenestration-inducing agent)^25^. In a recent study from 2025, Mao et al. showed that LSECs stained with CellMask Green and imaged using 3D SR-SIM exhibit signs of phototoxicity and reduced fenestration dynamics. Additional oxLDL treatment amplified the fenestration arrest by further increasing oxidative stress (in addition to photoactivated fluorophores)^16^.

In our report, by systemically screening fluorophores and testing ROS scavengers, we established conditions that preserved fenestration mobility, enabling the first long-term fluorescence imaging at a resolution beyond the diffraction of visible light. In particular, we demonstrated that fenestration dynamics can be monitored for over an hour without inducing cell toxicity or fenestration arrest. We conclude that 3D SR-SIM, in parallel to well-established AFM imaging, can be used for tracking the dynamics of fenestrations in live LSEC. Moreover, 3D SR-SIM allows overcoming the limitations of other techniques such as restricted field of view, low imaging speed or restricted selection of fluorophores.

Due to liver physiology, LSECs thrive in low oxygen conditions^26^. Still, the excessive depletion of oxygen was shown to disrupt many cellular processes and paradoxically promote ROS production in mitochondria^27,28^. Previous studies have shown that disruption of redox-regulating proteins, such as PDIA1, reduces fenestration number, supporting the idea that redox homeostasis directly influences actin-binding proteins and thus fenestration structure^29^. These findings explain the distinct effect of Oxyrase and NAC in our experiments. While NAC functions as a precursor to GSH, which in turn acts as a molecular ROS scavenger protecting intracellular structures^30–32^, other commercially available products focus on preventing ROS formation by scavenging oxygen. The latest is widely applied for fixed samples^33^, although compounds such as Oxyrase were also successfully applied in live cell imaging^34^. In this study, Oxyrase supplementation led to the formation of large gaps as well as a reduction in fenestration dynamics despite reducing the production of intracellular ROS. On the other hand, NAC not only reduced ROS formation and prevented phototoxicity-induced defenestration but also, by restoring the cellular redox balance, preserved fenestration dynamics without affecting the fenestration number. Overall, these results; as well as previously described mechanisms of NAC preventing _H2O2_-induced defenestration^12^, support the use of NAC supplementation for live cell 3D SR-SIM imaging.

Despite the fact that the exact molecular structure of LSEC fenestration remains unknown, we expect that ROS can disrupt both the fenestration-associated cytoskeleton^6^, membrane binding proteins^4,9^, cause membrane peroxidation^13^, as well as affect various cellular regulatory pathways^3,35^. In our previous work, we showed that H_2_O_2_ treatment increases intracellular ROS in LSECs and, in high concentrations, leads to severe defenestration and ultimately cell death^12^. Here, we reanalyzed our previously published datasets to quantify the fenestration speed parameter, which turned out to greatly resemble the phototoxicity resulting from 3D SR-SIM imaging presented in the current report. The effect of ROS on LSECs seems to be immediate and irreversible, causing first the loss of fenestration dynamics, then defenestration and in the later stage disruption of the whole cell membrane, resulting in cellular death. A similar effect on closing fenestrations was demonstrated using antimycin A. This mitochondrial complex III inhibitor generates a large amount of superoxide and leads to fenestration closing within 60 minutes. As ROS generation causes disruption in tubulin but no significant changes in the actin cytoskeleton (including actin forming fenestrae-associated cytoskeletal rings^6^), no immediate formation of stress fibers or cell thickening, it suggests that ROS most likely affect spectrin and other unknown proteins within the fenestration building complex^9,12^.

Our experiments concluded that only the combination of dye and illumination results in intracellular ROS formation, represented by strong DCF conversion in the mitochondrial region. Excited fluorophores can transfer excess energy to oxygen molecules, resulting in the formation of oxygen radicals and other reactive oxygen species. This process can lead to oxidative damage to various cellular components^36^. Previous reports showed that mitochondria are the primary site of ROS accumulation during live-cell imaging as well as suggested a significant differences in the amount of ROS formation between fluorophores^37–39^. The DCFDA/H2DCFDA probe, which we selected for our study, is not restricted to the detection of H_2_O_2_ and ROS, but also other one-electron-oxidizing species^40^, indicating a general increase in radicals after illumination. It was reported that increased intracellular glutathione (GSH) levels prevent DFCDA/H2DCFDA oxidation and improve cellular viability^41^. This suggests that NAC supplementation in our study helps LSECs to replenish GSH levels to mitigate phototoxicity. In addition, we observed that dyes prone to rapid photobleaching under 3D SR-SIM can also be used as a criterion for minimizing the ROS-induced phototoxicity during dye selection for live imaging. Our observation on LSEC nanomechanics demonstrated that significant cell softening (reduction in apparent Young’s modulus) was similar in cells exhibiting different labeling intensities in the same sample. This finding is especially important for the emerging studies proposing the use of live cell imaging with fluorescence-based methods for guiding the probe microscopy and spectroscopy^11,42^.

The proposed optimized imaging strategy addresses the fundamental challenge of live-cell imaging, therefore having a potential for broad applicability beyond fenestration research. Phototoxicity was limiting the use of optical techniques for e.g., live-cell studies of cytoskeletal dynamics, mitotic events, membrane dynamics and cellular transport^43–45^. In particular, the processes highly sensitive to oxidative stress, such as mitochondrial fusion/fission, endoplasmic reticulum remodeling or endocytosis will benefit from ROS scavenging^46–48^. There is a possibility that some existing live-cell optical studies have unknowingly generated and interpreted results affected by even subtle phototoxic artifacts. For example, membrane-associated protein NEMO involved in ‘active transport’ was later revealed as a phototoxic artifact^49^. Validation of existing findings using dedicated anti-phototoxicity protocols, such as NAC supplementation combined with optimized fluorophore selection, could provide new insights into cellular dynamics and increase awareness of this critical but often underestimated issue in live-cell microscopy^30–32^.

While the results presented here demonstrate that supplementation with NAC is a promising strategy to improve structural preservation during live-cell imaging through the regulation of intracellular ROS formation, further optimization can be implemented to enhance preservation of dynamic cellular processes. Those strategies may include: (i) engineering dedicated SIM environmental chambers with CO_2_ control for the flexibility in media selection; (ii) implementing total internal reflection fluorescence microscopy to provide lower energy transfer to restricted cell volume, thereby significantly reducing phototoxicity; (iii) targeting specific currently unknown fenestration-associated proteins, when those are identified; (iv) finding robust actin binding fluorophores to utilize indirect fenestration observation.

## 4. Conclusions

To our knowledge, this study represents the first reliable application of an optical imaging approach capable of resolving the dynamic behavior of liver fenestrations while avoiding previously reported artefacts. By combining careful fluorophore screening with ROS scavenging by NAC supplementation, we achieved hour-long 3D SR-SIM recordings that preserved both cell viability and fenestration mobility. Importantly, the combination of 3D SR-SIM with AFM provided complementary mechanistic insights into ROS-mediated phototoxic effect on fenestrations and LSEC nanomechanical properties. Moreover, our study highlights the challenges of optical live-cell imaging and establishes a framework that supports extended super-resolution imaging of cellular dynamics at the nanoscale, offering broad applicability well beyond fenestration research.

## 5. Material and Methods

### 5.1 Animals

For cell viability tests and 3D SR-SIM experiments, including ROS detection, RjHan:NMRI mice an in-house breeding line from the Animal Facility, faculty of Biology at Bielefeld University were used. For AFM and SEM, LSEC were isolated from wild-type male C57BL/6 mice. Mice were kept under standard conditions with water and chow (SSniff, regular chow diet) ad libitum. The mice used for the experiments were between 9 and 22 months old (SIM) and 4 – 9 months old (AFM). Animals were euthanized by cervical dislocation. All animal experiments were approved by the respective local authorities and were in accordance with institutional guidelines for the welfare of animals.

### 5.2 LSEC isolation and cell culture

Primary mouse LSECs were isolated as previously described in detail by Elvevold et al.^50^. In brief, the mouse livers were perfused and then enzymatically digested using 1.2 mg/50 mL Liberase (Roche). The non-parenchymal cell fraction is separated from the parenchymal cells by several centrifugation steps, followed by immunomagnetic separation of LSECs from the non-parenchymal cell fraction using CD-146 beads (MACS, Miltenyi). The cells were seeded according to the experimental design format described below.

### 5.3 Cell Viability assay

Cell viability was assessed by measuring the ATP using CellTiter-Glo 2.0 (Promega) assay performed according to the manufacturer’s guidelines. Isolated LSECs were seeded on 48-well plates coated with human fibronectin 0.2 mg/mL at a density of 2.5 × 10^5^ cells per well in Endothelial Cell Media (EGM) (Cell Applications, Inc.) containing 2% fetal bovine serum (FBS) overnight prior to the experiment. Cells were cultured at 37°C, 5% CO_2_, 5% O_2_ and 95% humidity. LSECs were treated with N-Acetylcysteine (NAC) (Sigma Aldrich) in varying concentrations (0.5, 1.0, 2.0 mg/mL) or with 1% Oxyrase (Sigma) for 1h. After the treatment duration, the cells were lysed and the Luminescence was subsequently measured using a plate reader (Tecan).

### 5.4 Live Cell Imaging

#### 5.4.1 Atomic force microscopy

LSECs were seeded on the bottom surface of plastic Petri dishes (TPP, Genos, Lodz, Poland) covered with human fibronectin (0.2 mg/mL) at a density of 65 000 cells in EGM-2 medium (Lonza, Basel, Switzerland) containing 2% fetal bovine serum (FBS) for 4 – 16 hours before measurements. Cells were cultured at 37°C, 5% CO_2_ and 95% humidity. The measurements were conducted using an atomic force microscope (AFM, Nanowizard IV, JPK Instruments/Bruker) at 37°C set using PetriDish heater™ (JPK Instruments/Bruker). SCM-PIC-V2 (Bruker) cantilevers (k = 0.1 N/m, nominal tip apex radius of 25 nm) were used for imaging in Quantitative Imaging (QI) mode, according to the methodology previously described for LSEC fenestrations^5,10^. Briefly, each image was acquired by performing multiple force curves in each pixel/point of the image that were translated into the images of topography and stiffness, where stiffness served only as a high contrasted image allowing detection of glass substrate (stiff – bright) and cell (soft – dark) and for evaluating the sharpness of the AFM probe. Load force was adjusted for individual cantilevers to achieve the best spatial resolution without distortion of fenestrations and was in the range of 200-350 pN. The length of the force curves (the *z* range) and the acquisition speed were in the range of 950 – 1250 nm and 100 – 140 µm/s, respectively. Consecutive images of the same area were used to create videos, allowing for the quantification of the dynamics of fenestrations. Additionally, pre-calibrated V-shaped cantilevers with the nominal spring constant of 0.03 N/m and the hemispherical tip (r = 5 µm) were used (MLCT-SPH-5UM-DC, Bruker) to assess the apparent Young’s modulus of cells, similar to the previous reports^8,9^. Before measurements, the AFM detector sensitivity was calibrated using a Petri dish surface without cells. For each experimental condition, 10-20 cells were measured using Force-Volume mode in the nuclear area (5 × 5 μm^2^). For each cell, 25 force-distance curves were acquired (loading force: 4 nN, loading rate: 8 µm/s, z range: 5 – 6 µm).

#### 5.4.2 Three-dimensional super resolution microscopy – structured illumination microscopy (3D SR-SIM)

Super-resolved fluorescence microscopy images were obtained using a 60× magnification objective lens with a numerical aperture (NA) of 1.42 on a DeltaVision OMX v4 3D super resolution-structured illumination microscope (Cytiva, Marlborough, MA, USA) (3D SR-SIM). 3D SR-SIM necessitates the acquisition of 15 images per z-plane, along with a minimum of six z-planes with a 125 nm distance, resulting in a total of 90 widefield illuminations. To ensure the inclusion of the fenestrations containing surface, we determined the requirement of recording a 1.5 µm thick z-stack, which translates to a total of 180 sample illuminations.

For all live cell imaging experiments, freshly isolated LSECs were seeded in EGM medium containing 2% FBS on round glass coverslips (Roth, 2.5 cm diameter) coated with human fibronectin (0.2 mg/mL) at a density of 1.5 × 10^6^ cells per well. Coverslips were incubated overnight at 37°C in 5% CO_2_, 5% O_2_, and 95% humidity before each experiment. During the live-imaging process, the coverslips were placed in a specially developed incubator, later called the “3D SR-SIM incubator”, which is adapted to the microscope stage and enables temperature monitoring during imaging.

#### 5.4.3 Experiment setup for evaluation of the effect of dye and illumination

Before measurements, the culture medium (EGM-2) was exchanged with 1.5 mL of fresh CO_2_ Independent Medium (Gibco™, ThermoFisher) (later called “imaging medium”) and placed on the inverted optical microscope connected with AFM. AFM was navigated using an inverted optical microscope (brightfield) to find the area of interest. Next, an area with fenestrations was selected, and 10 images in the same region were acquired. In the next step, the AFM head was removed, the light switched off, and the medium was replaced with the imaging medium containing fluorescent dye (7:1000, BioTracker). The AFM head was placed back on the sample and the measurement continued in the dark. After 25 – 30 minutes and collecting 10 scans of the same area, the AFM head was removed and the dye was rinsed ×3 with fresh imaging medium, and the AFM imaging was continued for 30 minutes. Finally, the illumination using a fluorescence lamp was turned on for 30 seconds to activate the dye, with subsequent scanning using AFM. The AFM scanning was continued for at least 30 minutes. In some experiments, the second event of illumination was introduced.

##### 5.4.3.1 Fluorescent dye testing

Commercially available live imaging-compatible dyes were tested on primary LSECs. The dyes included CellMask Green Actin Tracking Stain (Invitrogen, Thermo Fisher), CellMask Plasma Membrane Stain Orange (Invitrogen, Thermo Fisher), Vybrant DiI cell-labeling solution (Invitrogen, Thermo Fisher) and BioTracker 555 Orange Cytoplasmic Membrane Dye (Sigma). Each dye was tested individually. Cells were stained according to the manufacturers’ protocols followed by three rinses with fresh imaging medium. Stained cells on glass coverslips were placed into the 3D SR-SIM incubator. For each dye, a minimum of 20 cells were selected, and images were taken every 15 min over a total period of 60 min, using the respective excitation/emission wavelengths for each dye.

##### 5.4.3.2 Oxygen Scavengers/Antioxidants

To test the influence of the oxygen scavenger Oxyrase (Sigma) and the antioxidant NAC (Sigma) during the live-cell imaging process, LSECs were stained with BioTracker (Sigma) according to the manufacturer’s protocol. After staining, cells were washed three times with imaging medium and placed into the 3D SR-SIM incubator. Live-cell imaging was performed over a period of 45 minutes for 1% Oxyrase and up to one hour for 1 mg/mL NAC. For each supplement tested, cells from three different mice were analyzed, and at least five cells per treatment were evaluated. A control sample without any supplements was included for each treatment condition. Images were acquired every 15 minutes using an excitation wavelength of 568 nm.

##### 5.4.3.3 ROS Detection

For the detection of intracellular reactive oxygen species a ROS indicator (DCFDA / H2DCFDA - Cellular ROS Assay Kit, Abcam) was used according to the manufacturer’s protocol for fluorescent microscopy. The influence of illumination, fluorescent dye and oxygen scavengers NAC and Oxyrase on intracellular ROS amounts during live cell imaging was tested. LSECs were first stained with the fluorescent dye BioTracker according to manufacturer guidelines. Afterwards, the cells were washed ×3 with fresh imaging medium. To generate the normal value of intracellular ROS, the cells were incubated with the diluted DCFDA solution for 45 minutes at 37°C in the dark. Subsequently, an image was taken, given the starting point/normal value of ROS, which is referred to as “before illumination”. Again, diluted DCFDA solution was given to the cells, the same selected cells were then illuminated after every 3 minutes with an excitation of 568 nm for 45 minutes. The final image collected after 45 minutes is presented as “after illumination”. In the same way, the influence of the oxygen scavengers, NAC at a concentration of 1 mg/mL, and 1% Oxyrase during live cell imaging was tested. To test the influence of the dye itself, cells were also stained with the DCFDA solution and illuminated without prior staining with BioTracker. All images were recorded with the same laser intensity. To ensure linearity in the measurement of fluorescence intensity, the raw image data were analyzed using ImageJ/Fiji ^51^. Fold change was calculated by dividing the sum of fluorescence intensity after illumination by the intensity before illumination. For each condition, a minimum of 10 cells had been analyzed.

##### 5.4.3.4 Analysis of Fenestrations on 3D SR-SIM data

Analysis of fenestration count using ImageJ/Fiji^51^ and ilastik^52^ has been described elsewhere^53^. Briefly, 4D (XYZ and T) data stacks (DeltaVision files) were separated into individual files (corresponding to time point) via maximum intensity Z-projection. Pixel classification was done in ilastik, where representative images were used to indicate the areas of fenestrations and the area of the cell membrane. Afterwards, binary segmentation masks were created using batch processing and then analyzed using ImageJ/Fiji. First, the same thresholding value was applied to reduce bias. Then, binary masks were created and analyzed using “Analyze Particles” with parameters for size (>2500 – 16500 pixel) and circularity (0.4 – 1.0) for fenestrations.

##### 5.4.3.5 Analysis of dynamics of fenestrations

Analysis of fenestration mobility was performed using Hiro software (courtesy of the Department of Cell Biology at the Faculty of Biochemistry, Biophysics, and Biotechnology of the Jagiellonian University, Krakow, Poland)^54^. This program allows for the analysis of the change of position relative to the starting point of cells or cellular structures. Cell images obtained using AFM or SIM were used for analysis. These images were subjected to time-lapse analysis, in which the changes in the position of a single fenestration were tracked. Observed cell trajectories are presented as circular diagrams, in which the starting points of the trajectories are reduced to a common origin of the coordinate system.

## Acknowledgements

The authors gratefully acknowledge Prof. Peter McCourt, UiT, for linguistic revision of the manuscript.

