## Supplementary information for "Offsetting ROS-mediated arrest of endothelial fenestration dynamics permits long-term optical super-resolution- and AFM-imaging"

Table of content:

1. Additional examples of primary LSECs staining
2. Additional examples of LSEC fenestration dynamics during 3D SR-SIM live-cell imaging
3. Analysis of Fenestration Count in LSECs supplemented with 1% Oxyrase
4. Additional examples of LSECs investigated using AFM
5. LSEC nanomechanics

1. Additional examples of primary LSECs staining

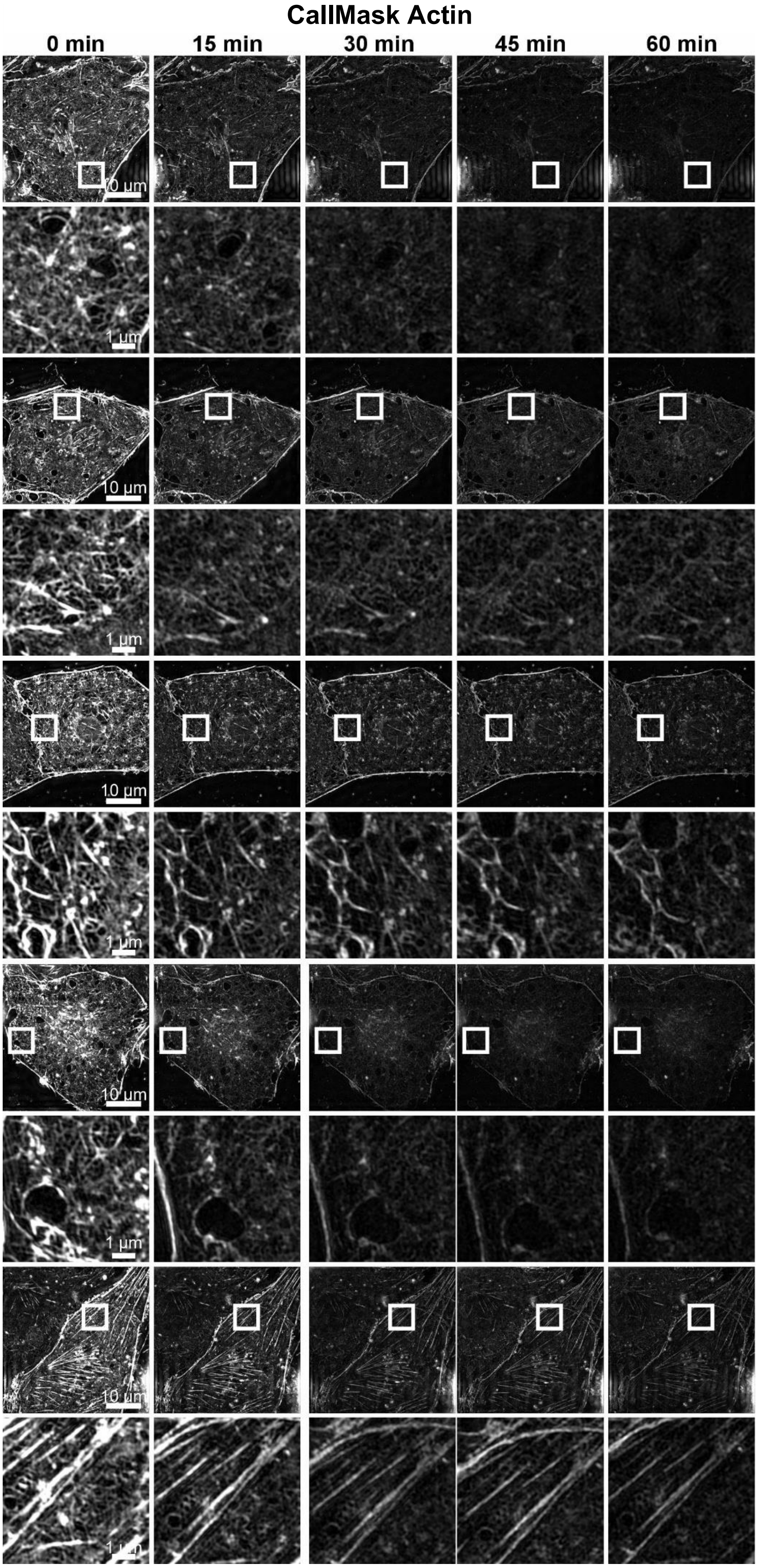


***Figure S1. Additional examples of primary LSECs stained with CellMask Green Actin Tracking Stain Orange.*** *Time-lapse 3D SR-SIM imaging of primary LSEC over one hour with 15-minutes intervals. Overview and ROI images demonstrate actin-associated labeling. The dye shows pronounced photobleaching, with a marked loss of fluorescence signal already after 15 minutes (two illuminations). These results further support the findings presented in* ***Figure 1****.*

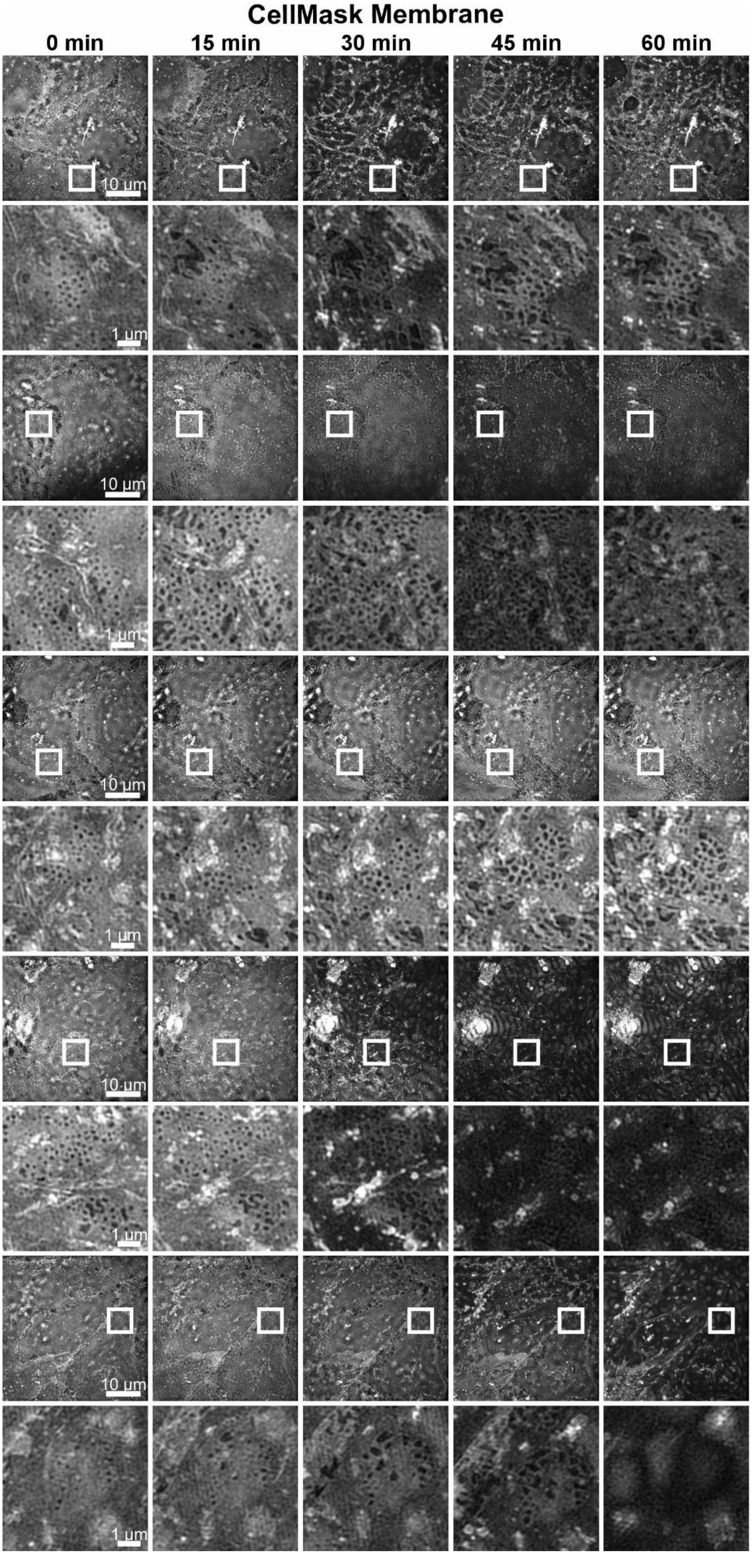


***Figure S2. Additional example of LSECs stained with CellMask Plasma Membrane Stain Orange***. *Representative time-lapse 3D SR-SIM images acquired over one hour at 15-minutes intervals. Overview and ROI images show a rather consistent membrane staining but with visible photobleaching in most cells after 15 minutes (2 illuminations). These findings further substantiate the results presented in* ***Figure 1***.

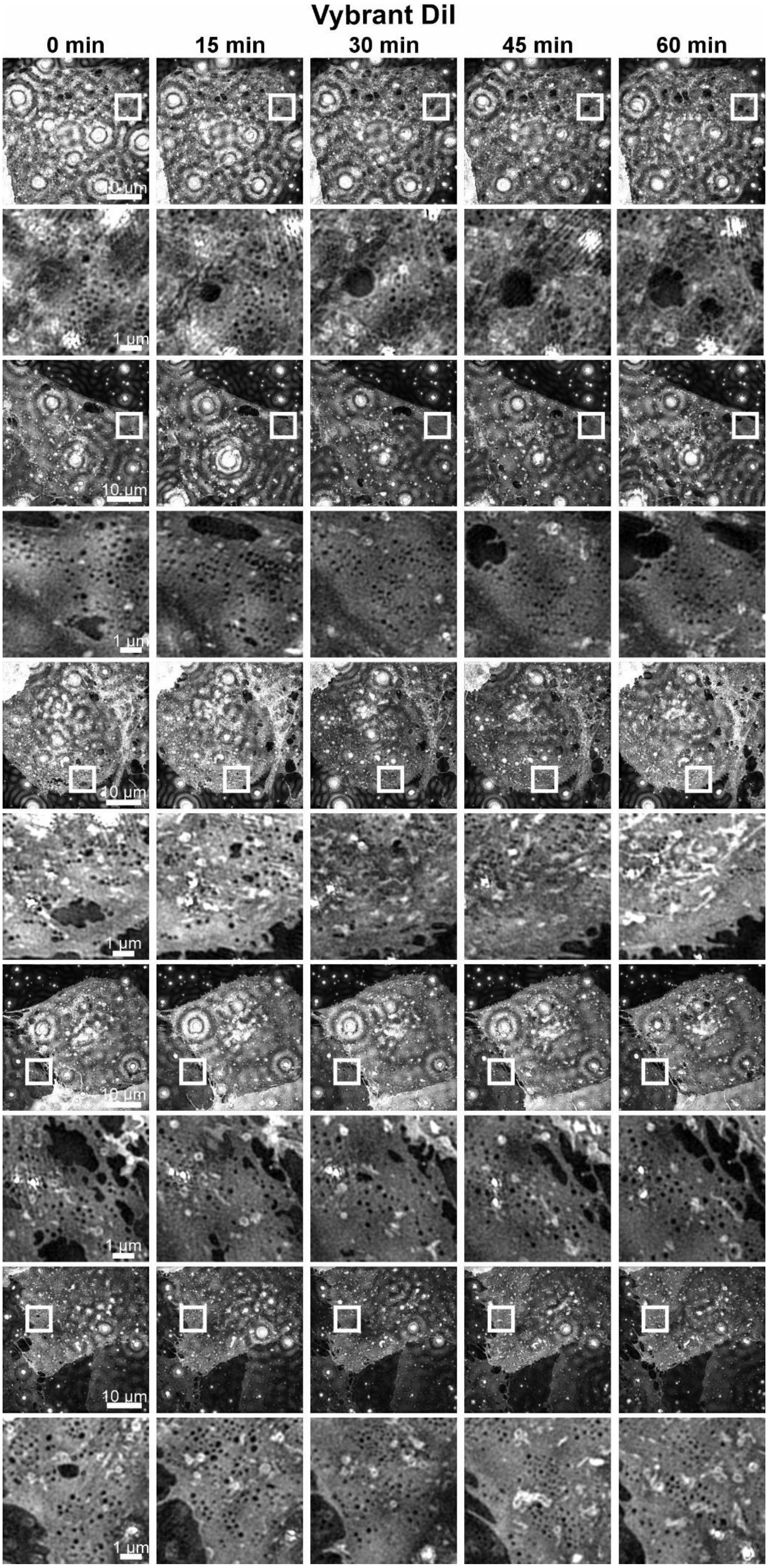


***Figure S3. Additional examples of LSECs stained with Vybrant DiI Cell-Labeling Solution.*** *Time-lapse 3D SR-SIM imaging of a primary LSEC over a one-hour period with 15-minute intervals. Overview and ROI images are provided to demonstrate dye performance and fenestration morphology. These data further illustrate the limited labeling quality and the consistent occurrence of bright dot-like artifacts as well as image reconstruction artifacts, which negatively interfere with the image quality presented in* ***Figure 1****.*

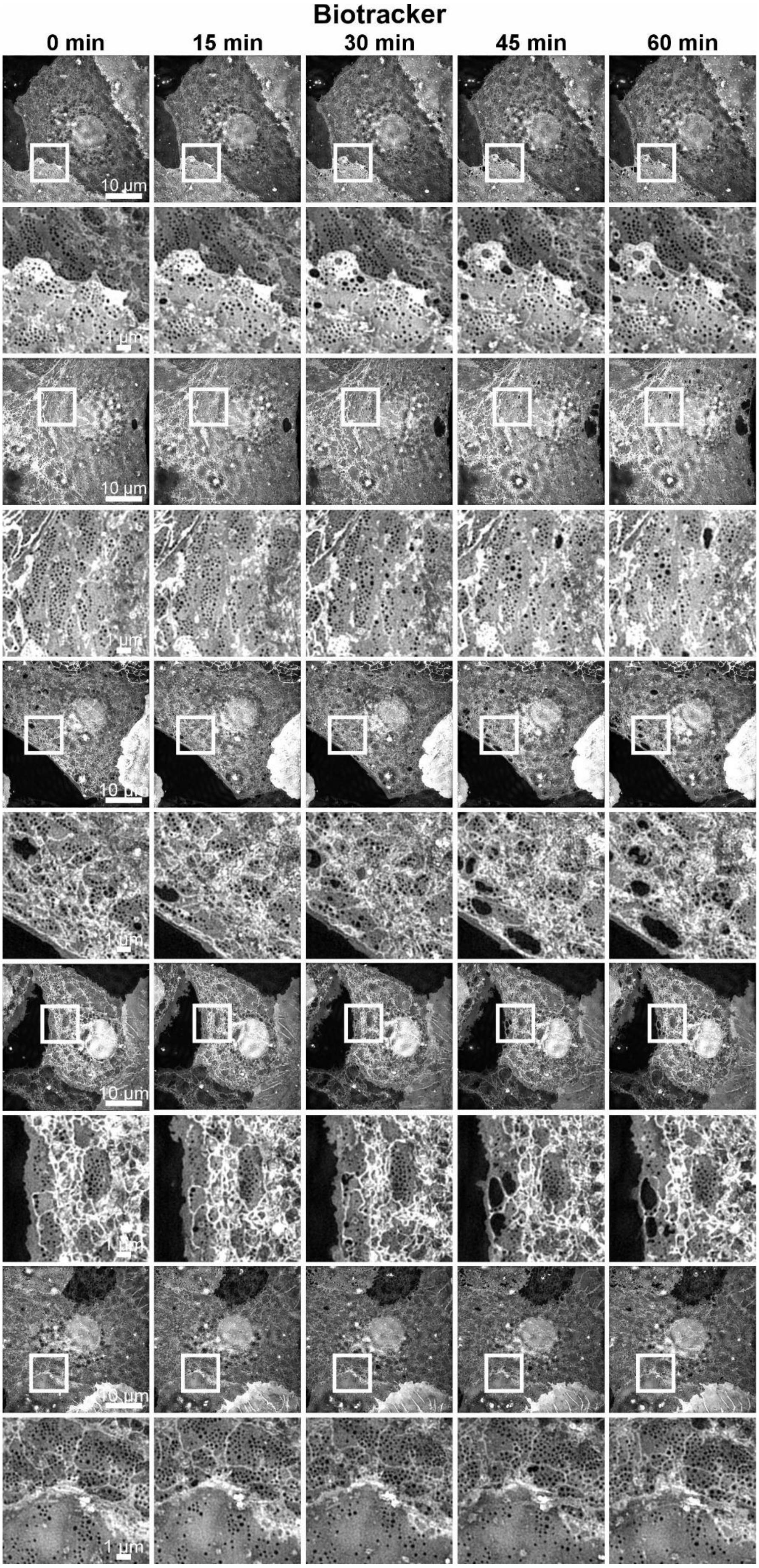


***Figure S4. Additional examples of LSECs stained with BioTracker 555 Orange Cytoplasmic Membrane Dye.*** *Representative time-lapse 3D SR-SIM images acquired over one hour at 15-minute intervals. Overview and ROI images show consistent membrane staining and clear visualization of fenestration morphology. This dataset supports the robust performance of BioTracker 555 during live-cell super resolution SIM, as described in* ***Figure 1****.*

2. Additional examples of LSEC fenestration dynamics during 3D SR-SIM live-cell imaging

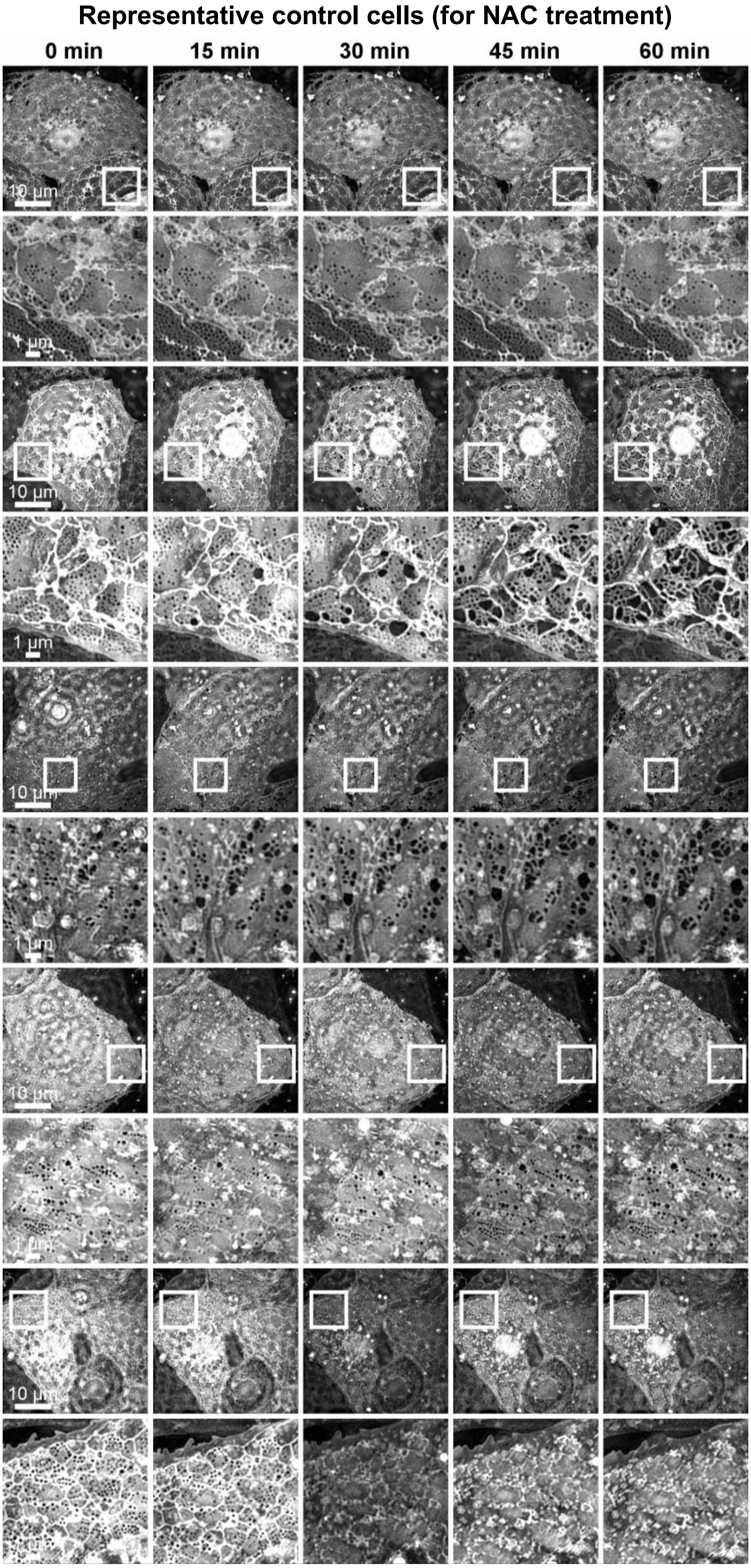


***Figure S5. Additional examples of LSEC fenestration dynamics during 3D SR-SIM live-cell imaging.*** *Time-lapse 3D SR-SIM images were acquired every 15 minutes over a 60-minute period. Shown are five additional representative cells under control conditions. Consistent with* ***Figure 3****, control cells exhibited arrested fenestrations with no detectable membrane remodeling, fenestration closure, or even membrane rupture.*

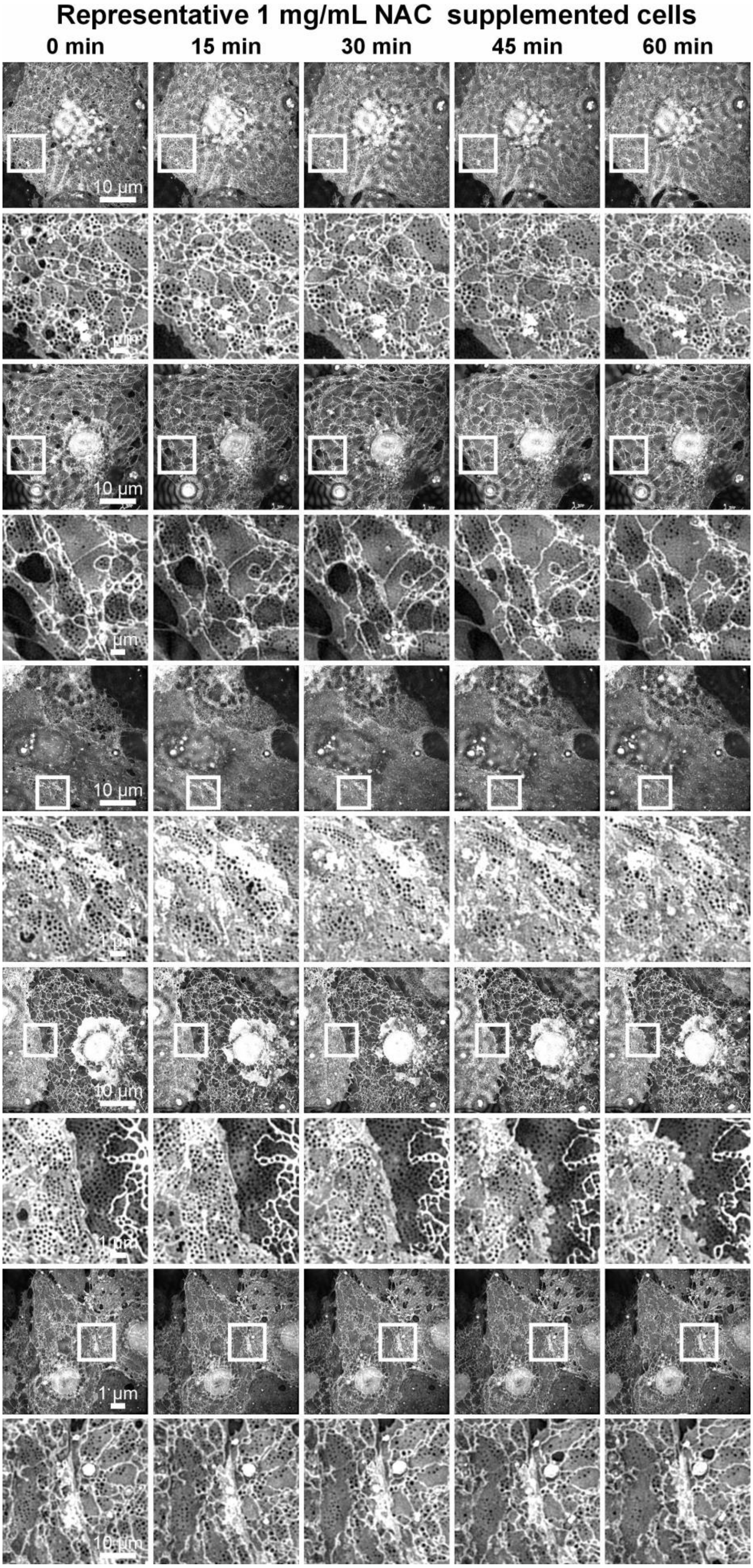


***Figure S6. Additional examples of the effects of NAC supplementation on LSEC fenestration dynamics during 3D SR-SIM live-cell imaging.*** *Time-lapse 3D SR-SIM images were acquired every 15 minutes over a 60-minute period. Shown are five additional representative cells with 1 mg/ml NAC supplementation. Consistent with* ***Figure 3****, NAC supplementation preserved fenestration mobility and active membrane dynamics.*

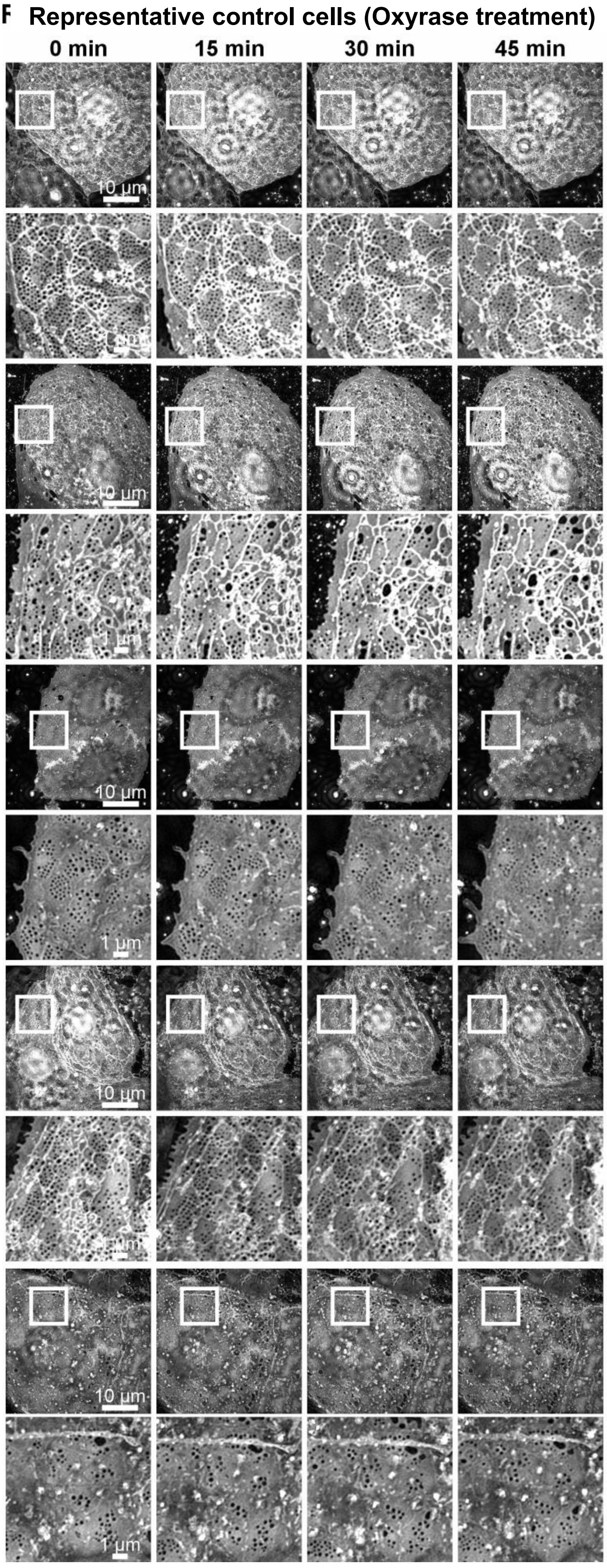


***Figure S7. Representative time-lapse 3D SR-SIM images of control cells without medium supplementation.*** *Time-lapse images were acquired over a 45-minute period, images were acquired every 15 minutes under identical conditions.*

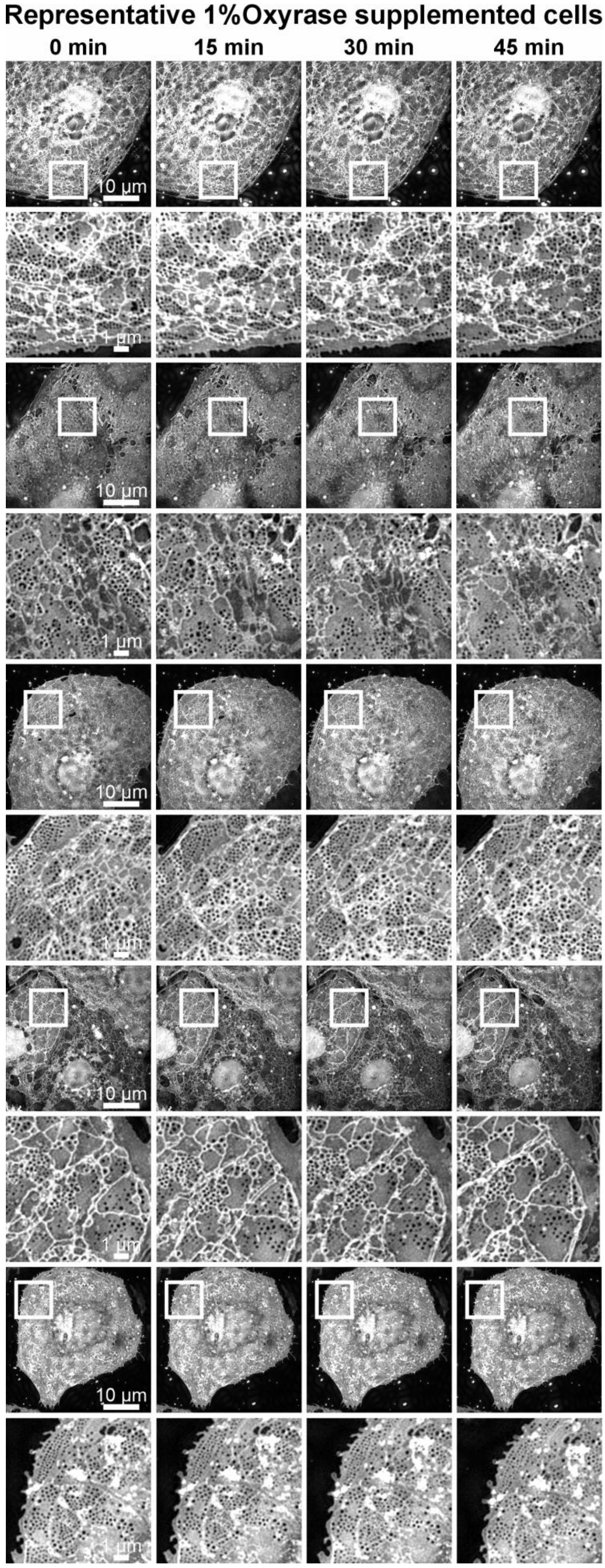


***Figure S8. Effects of Oxyrase supplementation on LSEC fenestration dynamics during 3D SR-SIM live-cell imaging.*** *Representative time-lapse 3D SR-SIM images of control cells with 1% Oxyrase medium supplementation. Time-lapse images were acquired over a 45-minute period, images were acquired every 15 minutes under identical conditions.*

### 3. Analysis of Fenestration Count in LSECs supplemented with 1% Oxyrase

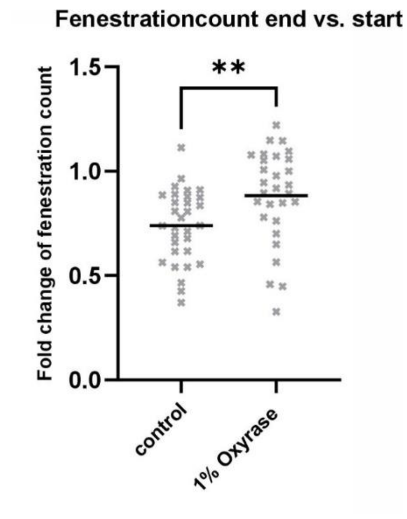


***Figure S9. Analysis of Fenestration Count in LSECs labelled with BioTracker and supplemented with 1% Oxyrase****. Quantification of fold change in fenestration count between the start and end of the experiment (60 min/0 min). No significant difference was detected between groups (Mean + datapoints; n = 3 animals, unpaired Mann–Whitney rank test; ** = p < 0.01). Each dot represents the same field of view tracked over the imaging period.*

### 4. Additional examples of LSECs investigated using AFM

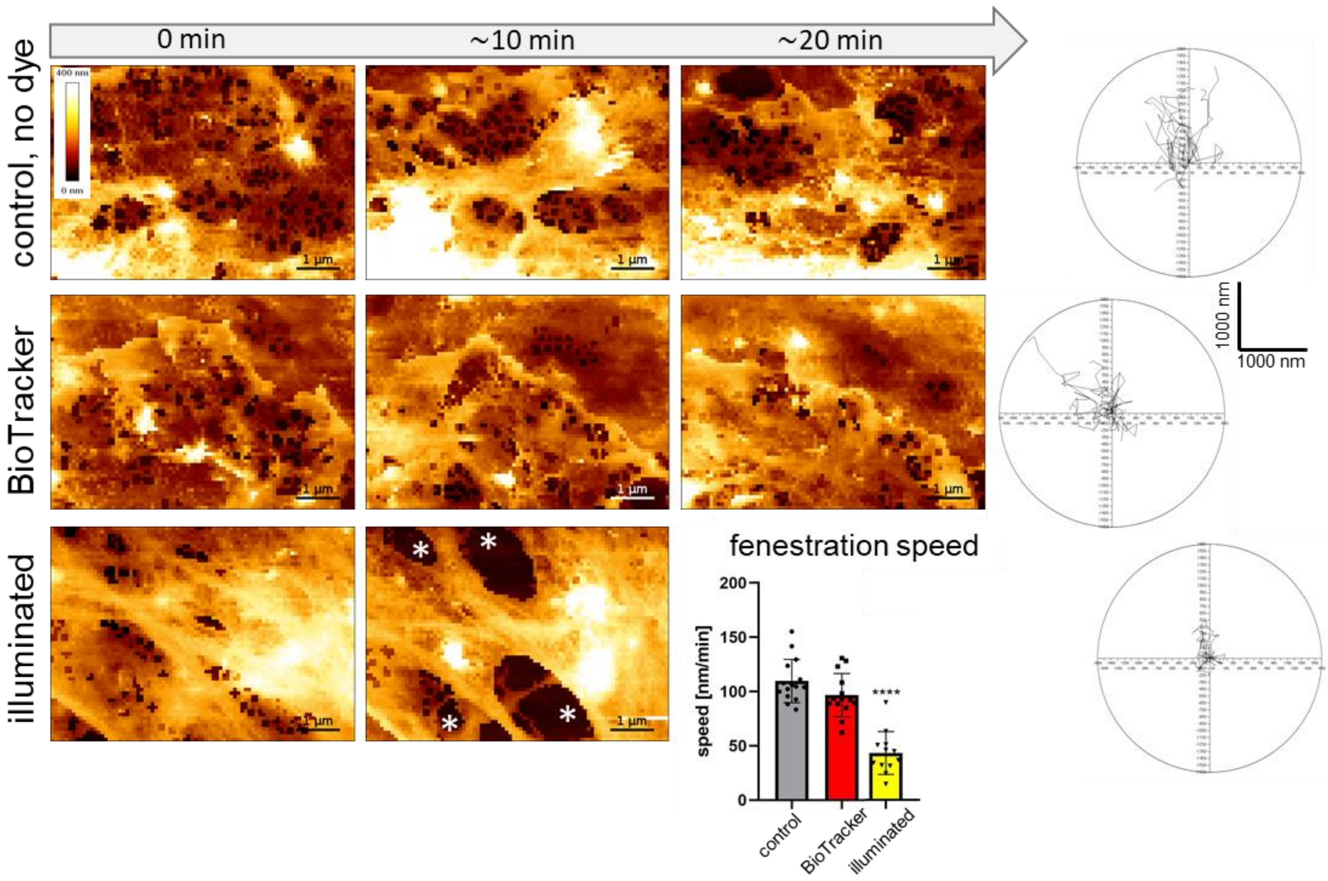


***Figure S10. Effects of 1% Oxyrase supplementation on LSEC fenestration dynamics during AFM live-cell imaging.*** *LSECs were measured in imaging medium at 37°C across all experimental stages. At stage 3, with the final rinsing, the imaging medium was supplemented with 1% Oxyrase. Stage 3 was limited to a few minutes and was not analyzed for fenestration trajectories to minimize the toxic effect of Oxyrase. Representative images are presented; remaining images are shown in* ***Video S3****. After Oxyrase, gap formation (asterisks) was observed during the illumination stage. Movement of remaining fenestrations was reduced by ~50%, as indicated in the circle plots of fenestrations trajectories and the chart.*

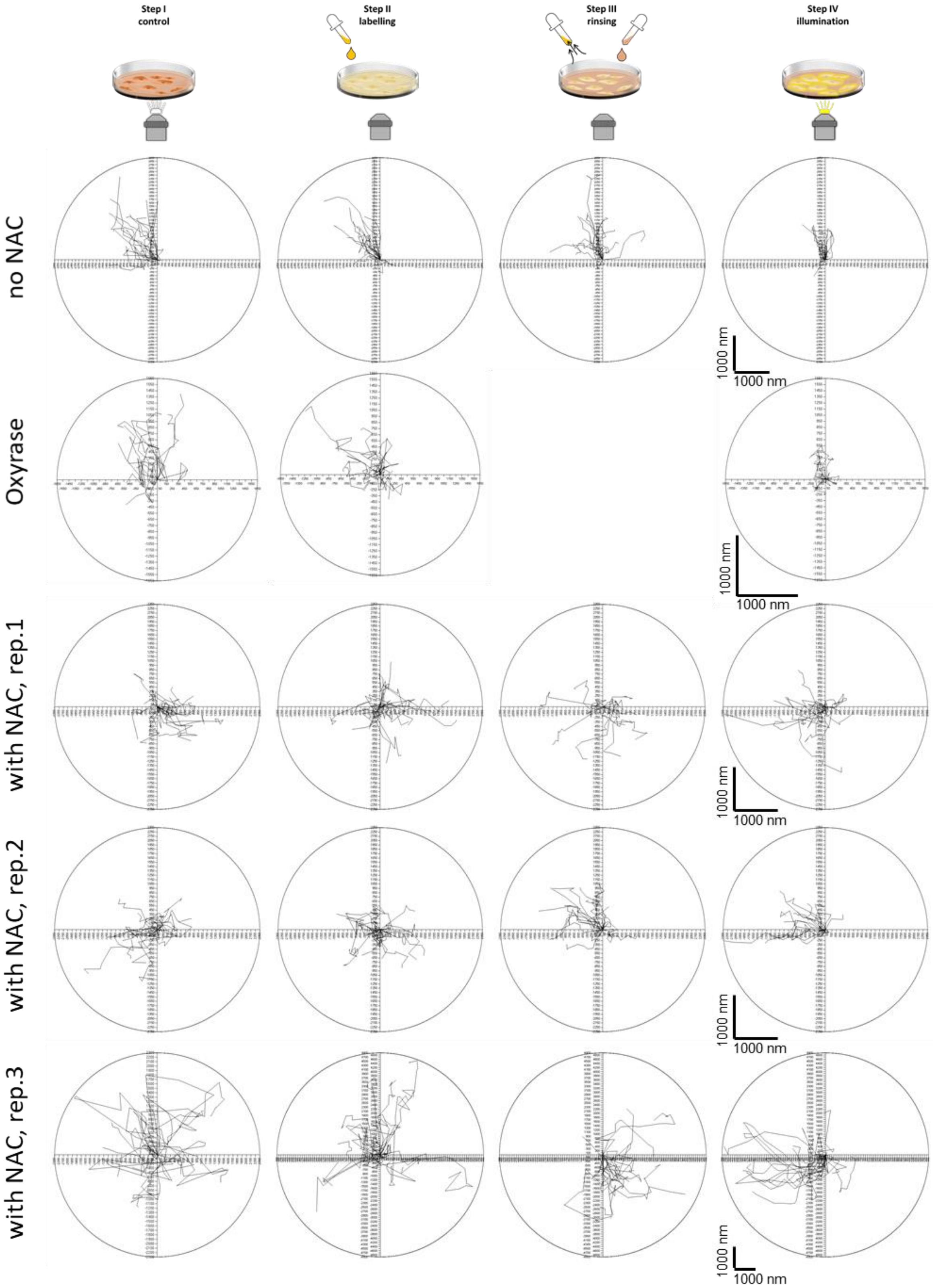


***Figure S11. AFM dynamics of LSECs labeled with BioTracker with or without NAC/Oxyrase supplementation.*** *Each plot represents the dynamics of individual fenestrations across the four stages of the AFM experiment: control, labelling with fluorescent dye, rinsing (not present for Oxyrase), and illumination (as described in Materials and Methods). Each trajectory in a chart corresponds to one fenestration tracked over time, enabling calculation of displacement and speed. The broad dispersion of trajectories indicates high fenestration dynamics. In the absence of NAC supplementation, fenestration displacement was markedly reduced, with movement largely restricted to collective shifts in the same direction.*

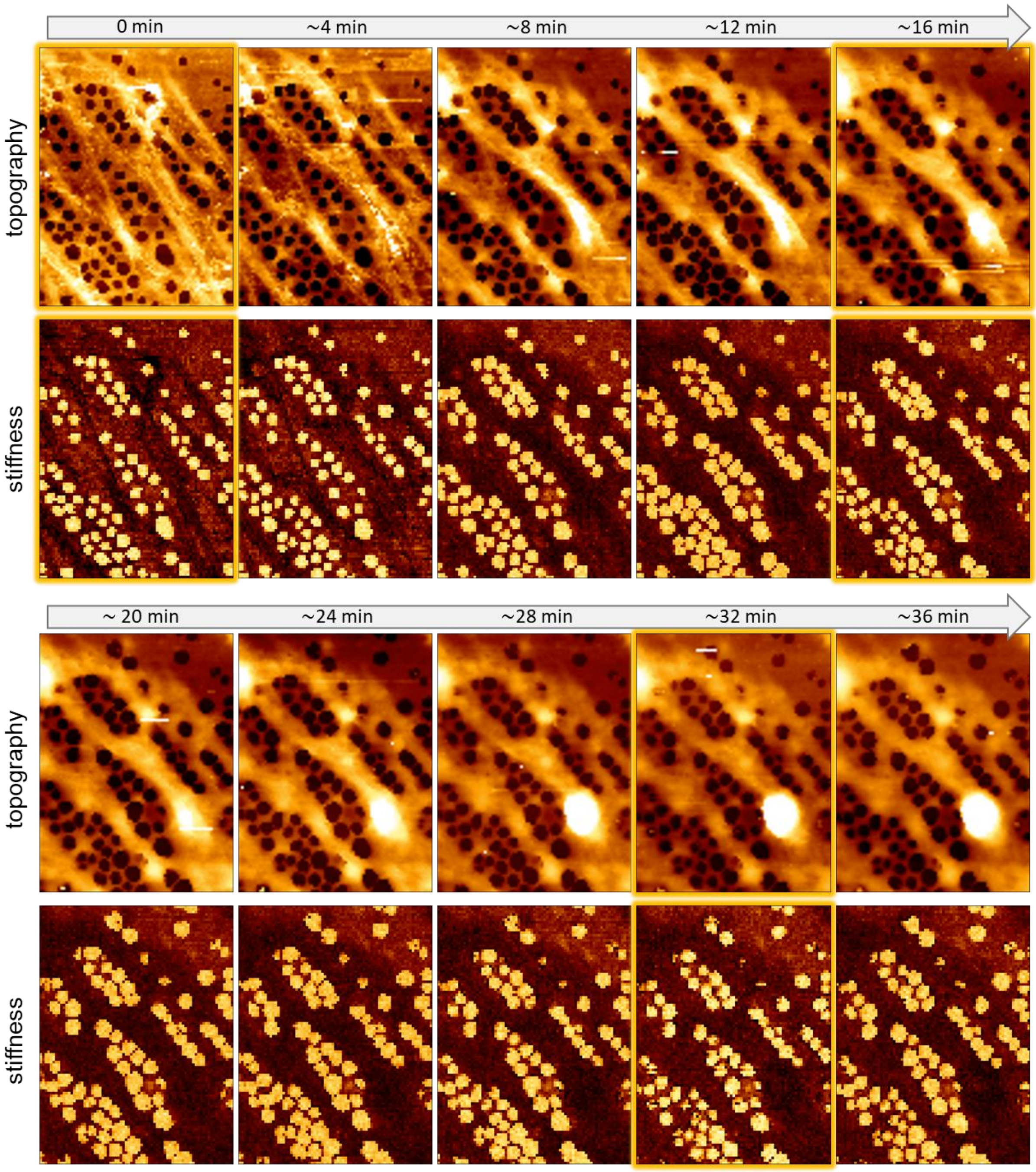


***Figure S12. Effects of CellMask membrane without supplementation on LSEC fenestration dynamics during AFM live-cell imaging.*** *LSECs were measured in imaging medium at 37°C across all experimental stages. Results starting from stage 3 are presented. Representative images show topography and stiffness (providing great contrast of fenestrations). When illuminated (images with gold glow effect) fenestrations diameters increased for a few minutes to shrink back to their original size. The test was performed three times and each time fenestration enlargement was observed. It should be highlighted that the dynamics of fenestrations is minimal and all fenestrations remain in the same position for the whole 38 minutes long experiment. The remaining images are shown in* ***Video S4****.*

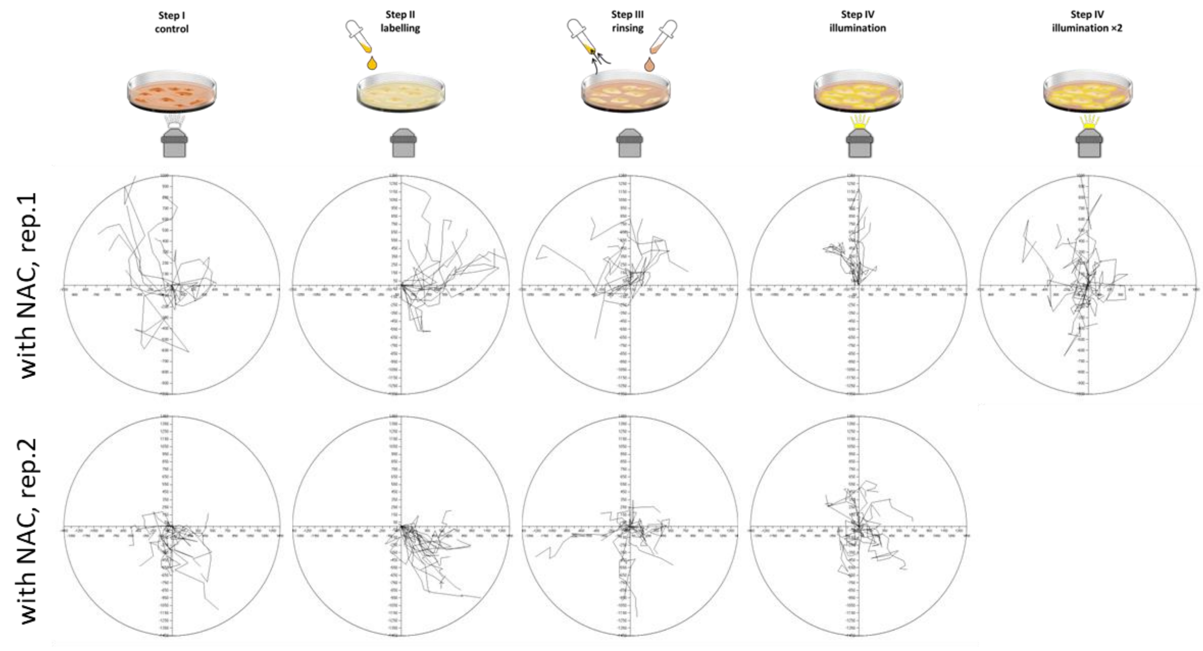


***Figure S13. AFM dynamics of LSECs labeled with CellMask actin dye in NAC-supplemented medium.*** *Each plot represents the dynamics of individual fenestrations across four stages of AFM experiment: control, labelling with fluorescent dye, rinsing, and illumination (as detailed in Materials and Methods). Each trajectory in a chart corresponds to one fenestration tracked over time, enabling calculation of displacement and speed. The broad dispersion of trajectories in all stages indicates high fenestration dynamics.*

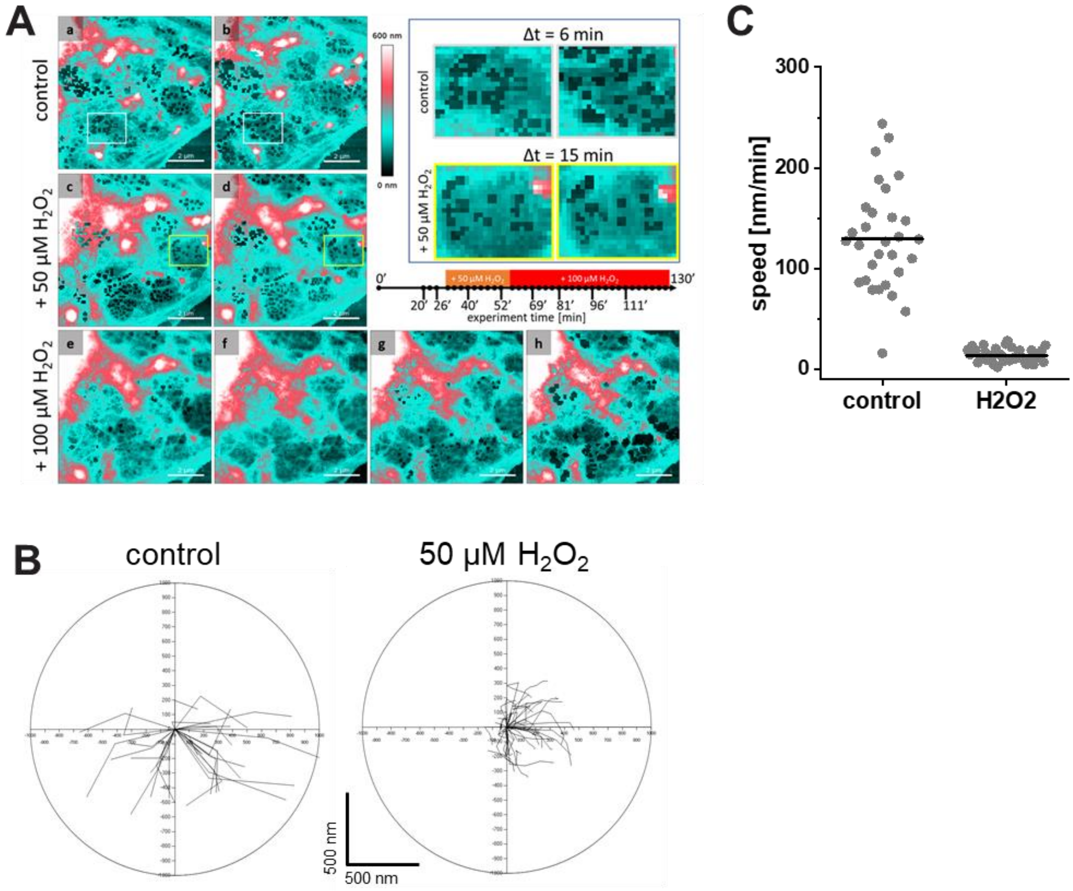


***Figure S14. ROS effect on the fenestration dynamics after 50 µM hydrogen peroxide treatment***. *A) Representative images of fenestrations at different time points (adapted from our previous study [12]).* ***B)*** *Fenestrations' positions were reanalyzed using Hiro software, as described in the Materials and Methods. Displacement of fenestration before and after treatment with 50 µM $H_2O_2$ was assessed. Δt was 6 min for the control and 24 min for the treated cells. Despite the time frame being four times shorter, the fenestrations migrated twice the distance of the treated group.* ***C)*** *Quantification of mean fenestration speed is shown for control and 50 µM $H_2O_2$-treated LSECs.*
